# Structural rearrangement of TFIIS- and TFIIE/TFIIF-like subunits in RNA polymerase I transcription complexes

**DOI:** 10.1101/353136

**Authors:** Lucas Tafur, Yashar Sadian, Rene Wetzel, Felix Weis, Christoph W. Müller

## Abstract

RNA polymerase (Pol) I is a 14-subunit enzyme that solely transcribes pre-ribosomal RNA. Cryo-EM structures of Pol I initiation and elongation complexes have given first insights into the molecular mechanisms of Pol I transcription. Here, we present cryo-electron microscopy structures of yeast Pol I elongation complexes (ECs) bound to the nucleotide analog GMPCPP at 3.2 to 3.4 Å resolution that provide additional insight into the functional interplay between the TFIIE/TFIIF-like A49-A34.5 heterodimer and the TFIIS-like subunit A12.2 present in Pol I. Strikingly, most of the nucleotide-bound ECs lack the A49-A34.5 heterodimer and adopt a Pol II-like conformation, in which the A12.2 C-terminal domain is bound in a previously unobserved position at the A135 surface. Our work suggests a regulatory mechanism of Pol I transcription where the association of the A49-A34.5 heterodimer to Pol I is regulated by subunit A12.2, thereby explaining *in vitro* biochemical and kinetic data.

## Introduction

Eukaryotic gene transcription is carried out by three different RNA polymerases (Pol). Pol I only transcribes pre-ribosomal RNA, which is subsequently cleaved into mature rRNAs in a complex process directly coupled to transcription (Schneider et al., 2007). In recent years, structural information available for *Saccharomyces cerevisiae* (yeast) Pol I has increased dramatically. Since the crystal structure of unbound Pol I was published in 2013 (Engel, Sainsbury, Cheung, Kostrewa, & Cramer, 2013; Fernandez-Tornero et al., 2013), cryo-EM structures of initiation (Engel et al., 2017; Engel, Plitzko, & Cramer, 2016; Han et al., 2017; Pilsl et al., 2016; Sadian et al., 2017) and elongation (Neyer et al., 2016; Tafur et al., 2016) complexes have become available. These structures revealed that while the Pol I initiation complexes greatly diverge from their Pol II (Plaschka et al., 2016) and Pol III (Abascal-Palacios, Ramsay, Beuron, Morris, & Vannini, 2018; Vorlander, Khatter, Wetzel, Hagen, & Muller, 2018) counterparts (suggesting a specific adaptation towards rDNA recruitment and transcription), the active Pol I enzyme adopts a conserved post-translocated conformation (in which the active site is free to bind the incoming NTP) as also observed in Pol II (Kettenberger, Armache, & Cramer, 2004) and Pol III (Hoffmann et al., 2015). In particular, in comparison with the apo Pol I crystal structure, the DNA-binding cleft is completely closed, the bridge helix adopts a fully folded conformation and the A12.2 C-terminal domain is excluded from the active site.

Compared to 12-subunit Pol II, 14-subunit Pol I and 17-subunit Pol III apparently incorporated transcription factor-like subunits during evolution (Khatter, Vorlander, & Muller, 2017; Vannini & Cramer, 2012). In Pol I, the A49-A34.5 heterodimer (hereafter referred as “heterodimer”) has been proposed to function as both a TFIIE- and TFIIF-like factor, participating during transcription initiation and elongation (Khatter et al., 2017; Vannini & Cramer, 2012). While the A49 C-terminal tandem winged helix domain (tWH) has structural homology to TFIIE, the N-terminal A49 dimerization domain, together with the A34.5 subunit, form a triple β-barrel structure that resembles the Rap74/30 module of TFIIF (Geiger et al., 2010). In accordance with its proposed TFIIE-like function, the A49 tWH has been observed bound to upstream DNA during initiation (Engel et al., 2017; Han et al., 2017) and in an elongation state with short RNA (Tafur et al., 2016). However, this domain is flexible and not seen when the transcription scaffold contains longer RNAs or in the absence of nucleic acids. The heterodimer is anchored to the core enzyme by interactions between the A49-A34.5 dimerization domain with the N-terminal domain of A12.2, the A135 lobe, and by an extended surface between the long C-terminal tail of A34.5 (A34.5-Ct) and the A135 surface (Engel et al., 2013; Fernandez-Tornero et al., 2013). The A34.5-Ct forms the largest interaction surface and (like A49) is essential for stable binding of the heterodimer to Pol I *in vivo* (Gadal et al., 1997) and *in vitro* (Geiger et al., 2010). Accordingly, Pol I purified from an *rpa34-*Δ yeast strain lacks A49 (Gadal et al., 1997), and that purified from a strain with a deletion of A49 lacks A34.5 (Pilsl et al., 2016).

Since its discovery, Pol I has been shown to exist in two different conformations that differ by the presence of the heterodimer, which can be reversibly dissociated(Huet, Buhler, Sentenac, & Fromageot, 1975). The form lacking A49-A34.5, termed Pol I*, has reduced transcriptional specificity and activity compared to the complete Pol I enzyme (Huet et al., 1976). Interestingly, Pol I* is also more sensitive to inhibition by α-amanitin (Huet et al., 1975). Neither A49 (Liljelund, Mariotte, Buhler, & Sentenac, 1992) nor A34.5 (Gadal et al., 1997) are essential genes, and Pol I* has been proposed to co-exist with Pol I *in vivo* (Gadal et al., 1997). Deletion of topoisomerase I causes a very strong growth defect in yeast only when combined with a deletion of A34.5 (Gadal et al., 1997), suggesting that A34.5 is important for relieving topological stress during rDNA transcription. *In vitro*, the heterodimer has a stronger effect on promoter-dependent transcription than on non-specific transcription, which depends entirely on the presence of the A49 tWH (Pilsl et al., 2016). When using a transcription bubble or a minimal nucleic acid scaffold (neither of which needs strand opening), the absence of heterodimer affects primarily the processivity of the enzyme (the ability to transcribe the entire transcript) (Geiger et al., 2010; Kuhn et al., 2007). Interestingly, in the mammalian system, the association of the PAF53-PAF49 heterodimer (the human ortholog of A49-A34.5) to the Pol I core is regulated by the nutrient status of the cell, whereby upon nutrient starvation the PAF53-PAF49 heterodimer dissociates from Pol I and is depleted from the nucleolus (Penrod, Rothblum, Cavanaugh, & Rothblum, 2015). Overall, the data suggests that the heterodimer is functionally important for transcription initiation and/or elongation. However, the functional and physiological relevance of Pol I* has not been elucidated to date.

A12.2 is another Pol I-specific subunit which shares homology with Pol II subunit Rpb9 (in its N-terminal domain) and Pol II transcription factor TFIIS (in its C-terminal domain). While the role of TFIIS in RNA cleavage is well established (Cheung & Cramer, 2011), Rpb9 appears to regulate transcription elongation (Hemming et al., 2000), proofreading (Knippa & Peterson, 2013) and transcription-coupled DNA repair (Li, Ding, Chen, Ruggiero, & Chen, 2006). The A12.2 C-terminal Zn ribbon domain (A12.2C) is required for the Pol I intrinsic RNA cleavage activity (Kuhn et al., 2007) and adopts a similar position as TFIIS in the cleft in unbound (apo) Pol I (Engel et al., 2013; Fernandez-Tornero et al., 2013; Neyer et al., 2016) (as well as in Pol I bound only to DNA (Sadian et al., 2017; Tafur et al., 2016)), but is excluded from the active site upon formation of the EC (Neyer et al., 2016; Tafur et al., 2016). Its exact position, however, has not been determined in the context of an actively transcribing complex. While deletion of the A12.2C does not cause any growth defect, deletion of the A12.2 N-terminal Zn ribbon domain (A12.2N) produces a similar effect as deletion of the complete protein (growth defect at 34°C, synthetic lethality when combined with A14 deletion, and strong sensitivity towards 6-azauracil and mycophenolate) (Van Mullem, Landrieux, Vandenhaute, & Thuriaux, 2002). Interestingly, deletion of either the complete A12.2 or A12.2N also alters the nucleolar localization of Pol I, suggesting that A12.2 is important for Pol I integrity.

Studies to date suggest a functional interplay between the Pol I heterodimer and subunit A12.2. The heterodimer stimulates A12.2-mediated RNA cleavage *in vitro* (Geiger et al., 2010), the latter which is important for Pol I backtrack recovery (Lisica et al., 2016). A12.2N interacts directly with the dimerization domain of A49, thus stabilizing the anchoring of the heterodimer (Engel et al., 2013; Fernandez-Tornero et al., 2013). Recently, A12.2 has also been proposed to be important for transcription initiation *in vivo* and *in vitro*, especially in the absence of A49 (Darrière T, 2018). Therefore, it is likely that both A49-A34.5 and A12.2 increase the processivity of the enzyme *in vivo.*

While the Pol I elongation complex (EC) greatly resembles Pol II and Pol III ECs, some elements in the active site appear to be positioned differently in Pol I, including two loops (loop A and loop B) from the A135 lobe that come close to the non-template (NT) single strand (ssNT) in the transcription bubble. Interestingly, even though most residues near the active site are conserved between the three Pols, mutations in some of these appear to have opposite effects in Pol I and Pol II *in vitro* (Viktorovskaya et al., 2013). These results suggest that there might be differences in the active site that could be observed in other EC intermediates or in a pre-translocated state, in which the nucleotide binding site (“A” site) is occupied by the incoming NTP substrate. Additionally, both the A12.2 subunit and the A49-A34.5 heterodimer might play active roles during elongation that have not been structurally characterized to date.

## Results

### Cryo-EM structures of the GMPCPP-bound Pol I elongation complex (EC)

In order to better understand the catalytic mechanism of Pol I, we incubated the Pol I EC with the non-hydrolysable NTP analog GMPCPP as previously used for Pol II (Kettenberger et al., 2004; Wang, Bushnell, Westover, Kaplan, & Kornberg, 2006). The Pol I EC was prepared as previously described (Tafur et al., 2016), except that 1 mM of MgCl_2_ was included in the buffer (Material and Methods). 5,768 micrograph movies were collected on a FEI Titan Krios equipped with a K2 direct electron detector, and processed with RELION 2.0 (Kimanius, Forsberg, Scheres, & Lindahl, 2016). After sorting particles with 2D and 3D classification, an unexplained extra density next to the A135 surface was observed in most of the particles with a closed cleft and strong DNA-RNA density, concomitant with streaky and weak density for the A49-A34.5 heterodimer. To better resolve this density, particles were classified using a mask in this region (Fig. 1-Figure Supplement 1). This revealed that the extra density corresponded to A12.2C (Fig. 1). In total 63% of all particles selected after the first unmasked 3D classification step did not have the heterodimer bound and showed density for A12.2C in this new position (named Pol I* (Huet et al., 1975)), while only 37% represented the 14-subunit Pol I. Extensive 3D classification ultimately yielded two different, nucleotide bound ECs (referred throughout the text simply as ECs): 12-subunit Pol I* EC lacking the heterodimer, which was refined to 3.18 Å resolution, and 14-subunit Pol I EC, which was refined to 3.42 Å resolution. The overall conformation of both Pol I forms is very similar, with the exception of the presence/absence of the heterodimer and a slight difference in the conformation of the clamp, and resembles previously published structures (Figure 1-Figure Supplement 3) (Neyer et al., 2016; Tafur et al., 2016).

**Figure 1.**
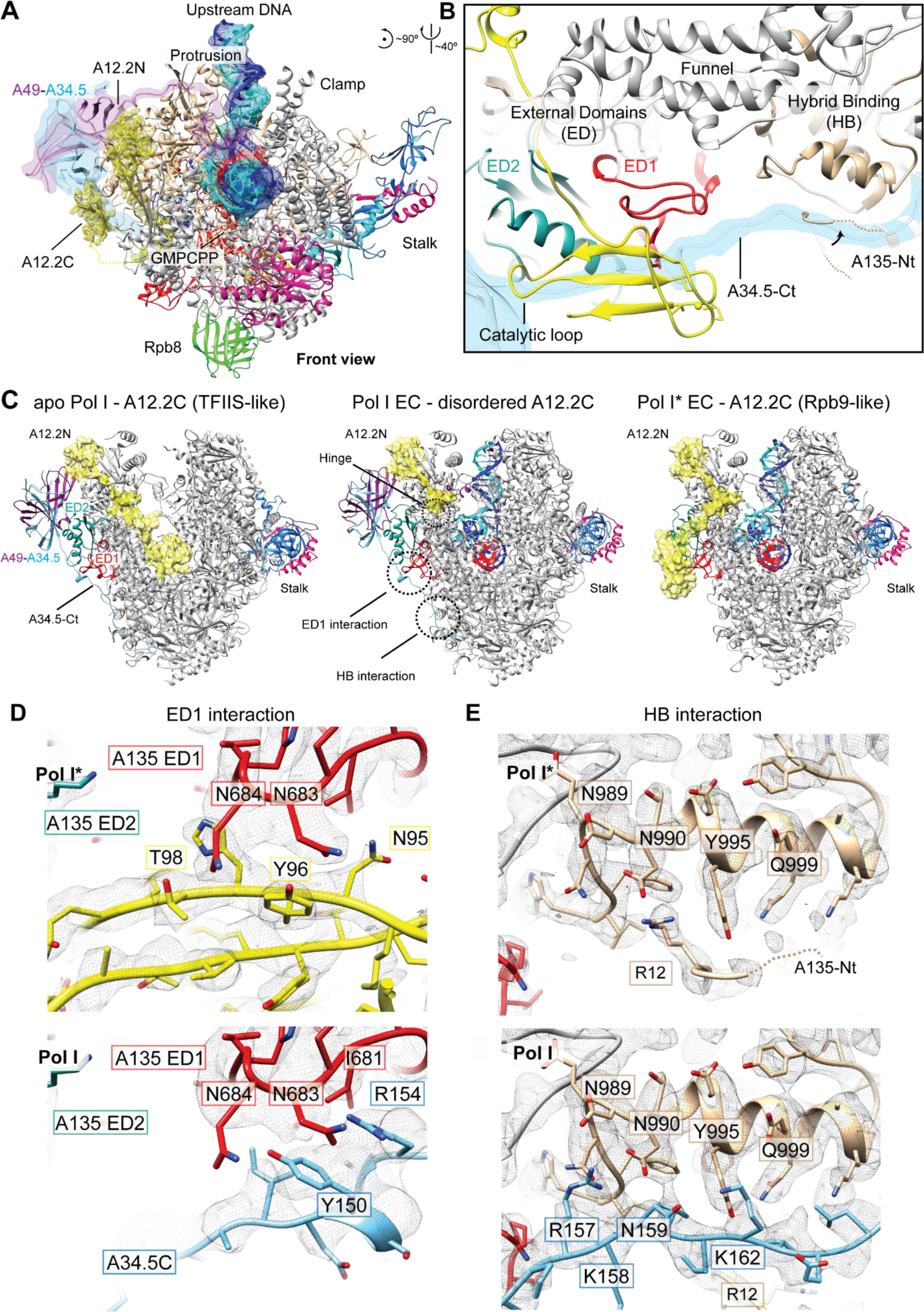
Structure of the Pol I EC bound to GMPCPP and structural rearrangement of the A12.2 C-terminal domain. **A.** In Pol I*, the A12.2 C-terminal domain (A12.2C) can bind to an Rpb9-like site next to the A135 ED1, which clashes with the position of the A49-A34.5 heterodimer in Pol I. Density for the DNA is from the Pol I (core) EC (upstream DNA) reconstruction (unsharpened), while the density for the A12.2C is from the Pol I* EC (+GMPCPP) map. **B.** Two interfaces are differently arranged in Pol I* versus Pol I. Both A34.5 and A12.2 can bind to the A135 External Domain 1 (ED1), and A34.5 and the N-terminal tail of A135 (A135-Nt) can bind to the Hybrid Binding domain (HB). **C.** Comparison between the apo (left), Pol I EC (middle) and Pol I* EC (right) reveals that the A12.2C can alternate between TFIIS-like (apo) or Rpb9-like (right) positions. Movement of the A12.2C is around a hinge at residue 67, indicated in the Pol I EC, where the A12.2C is disordered and not observed. The position of the ED1 and HB interaction surfaces are indicated in the Pol I EC. A12.2 is shown as ribbon diagram and yellow surface for easier visualization. **D.** Specific interactions of the ED1 with either A12.2C (Pol I*, top) or A34.5 (Pol I, bottom). **E.** Specific interactions of the HB with either A135-Nt (Pol I*, top) or A34.5 (Pol I, bottom). Densities shown for panels D and E are from the sharpened Pol I* and Pol I EC (+GMPCPP).

Interestingly, an apo Pol I* reconstruction at 3.21 Å resolution was obtained with a similar conformation as previously observed for the cryo-EM structures of monomeric Pol I (Neyer et al., 2016; Pilsl et al., 2016), highlighting that the presence of the heterodimer does not impose any conformational constraints on the Pol I core. In this reconstruction, the bridge helix is unfolded, the cleft is only partially closed and the DNA-mimicking loop is excluded from the active site (Fig. 1-Figure Supplement 4). At present it is unclear why most of the particles lack the heterodimer compared to previous Pol I EC structures (Neyer et al., 2016; Tafur et al., 2016), but it is possible that the presence of GMPCPP promoted the exclusion of the heterodimer, thus favoring the equilibrium towards the Pol I* EC in the sample.

Models were built using previous Pol I structures as a starting point and were real-space refined, yielding structures with excellent stereochemistry (Table 1). The structures obtained in this work reveal: (1) a previously unobserved position of A12.2C not compatible with A49-A34.5 heterodimer binding, (2) the position and interactions of the incoming NTP substrate in the active site, and (3) the stabilization of the downstream end of the transcription bubble by flipping of the +2 base and stacking of the +1 base of the non-template strand.

**Table 1.**
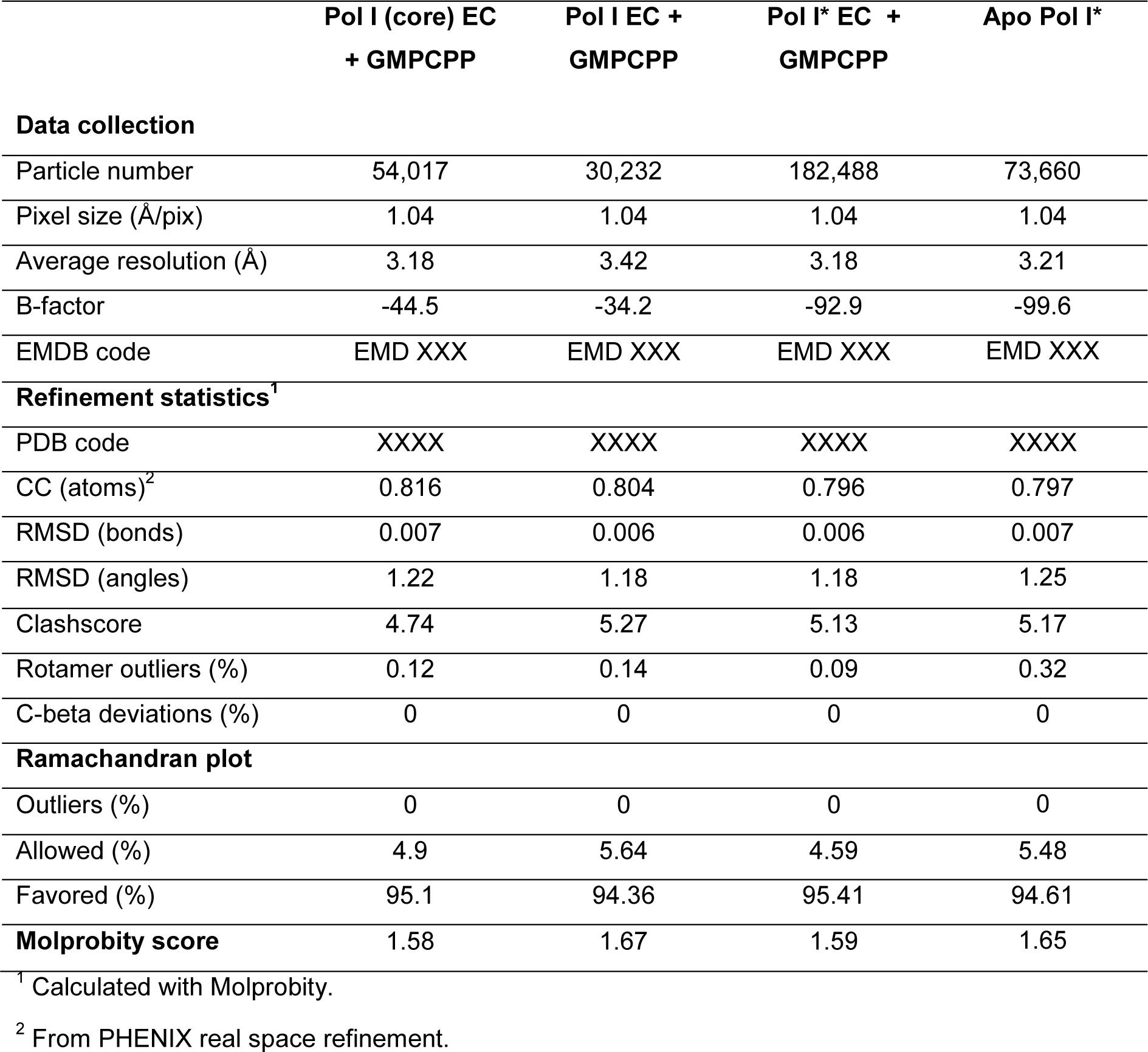
Data collection and refinement statistics.

### The A12.2 C-terminal domain alternates between a TFIIS-like and an Rpb9-like position

In the complete, 14-subunit Pol I EC, the A12.2C is disordered and only density up to residue 67 is observed, as previously described (Neyer et al., 2016; Tafur et al., 2016). In the Pol I* EC, however, the A12.2C is anchored to the enzyme by interactions with the A135 External Domain 1 (ED1) in a novel position that overlaps with the A34.5 C-terminal tail (A34.5-Ct) in the Pol I EC, and which is similar to the position of the C-terminal domain of Pol II subunit Rpb9 (Rpb9C) (Fig. 1A, see below). A12.2N and A12.2C are connected by a flexible linker that is weakly resolved in the Pol I* EC, although not at sufficient resolution for model building but further validating the assignment of this density (Figure 1-Figure Supplement 3). Interestingly, additional density is observed from the A190 jaw domain that connects to the DNA-mimicking loop (DML) towards the A12.2 linker, suggesting that the DML might also contribute to this positioning of the A12.2C (Figure 1-Figure Supplement 3).

Comparison of Pol I* with Pol I reveals that two interfaces in the A135 surface are differently arranged (Figure 1B). In addition to the interaction with the ED1, which mutually exclusive binds either A34.5-Ct or A12.2C, residues 989 to 1000 from the A135 Hybrid Binding domain (HB) interact with either the A34.5-Ct or the A135 N-terminal tail (A135-Nt), which folds back towards the HB in Pol I* (Figure 1B, C). The A135-Nt effectively acts as a switch, changing its positioning to allow or to prevent A34.5-Ct binding to the HB. Such a movement is only possible because the first helix of A135 is permanently associated with parts of AC40, A135 and Rpb10, and only the flexible tail (residues ~1-18) can move freely. The A12.2C can alternate between the previously observed TFIIS-like and the Rpb9-like position by rotating around a hinge located at residues 66-67, just after the well-ordered helix inserted between the funnel and jaw (A12 residues 59-65) (Fig. 1C, middle).

Both A12.2C and A34.5-Ct interact with two asparagine residues in the ED1 (A135 N683 and N684) through a tyrosine residue (A12.2 Y96; A34.5 Y150). A12.2C interacts with N683 and N684 through Y96, and with N684 through A12.2 T98 (Fig. 1D, top). On the other hand, A34.5 Y150 interacts with N684, and the neighboring R154 with N683 (Fig. 1D, bottom). A similar situation is observed in the HB interaction surface. In Pol I, A34.5 R157 is close to residues A135 N989 and N990 (Fig. 1E, bottom). In the Pol I*, R157 is partially replaced by A135 R12, which comes near residue N990 (Fig. 1E, top). Thus, two interaction surfaces of A34.5-Ct with A135 are exchanged in Pol I* with A12.2C (ED1) and the A135-Nt (HB), which interact with the same residues from A135. These interactions preclude the binding of the heterodimer when A12.2C adopts the Rpb9-like position.

In the monomeric apo Pol I, in which the cleft is partially closed, A12.2C can still occupy the TFIIS-like position (Neyer et al., 2016). However in the apo Pol I*, A12.2C is stably bound to the Rpb9-like position, although there is sufficient space for accommodating A12.2C in the TFIIS-like position (Fig. 1-Figure Supplement 4). The presence of the heterodimer in the enzyme could thus promote binding of A12.2C to the TFIIS-like site (when accessible) by blocking the Rpb9-like binding site. Comparison of apo Pol I with apo Pol I* reveals that the change in the position of A12.2C also shifts A12.2N by ~3 Å towards the jaw, and part of the latter appears to move towards the linker, likely to stabilize its position (Fig. 1-Figure Supplement 4). Interestingly, both domains move relative to the region that contributes the fifth β-sheet to the jaw (resides ~43-66), which fixes A12.2 to the Pol I core. Therefore, the movement of both, the A12.2N and the jaw, accommodate the change in the position of A12.2C.

### The structure of the ED1 determines binding of the C-terminal domain of the Rpb9-like subunit in Pol I, Pol II and Pol III

Comparison of the Pol I/Pol I* structures with Pol II and Pol III reveals that while the ED2 appears to be structurally more conserved, the Pol I ED1 diverges from its Pol II and Pol III counterpart, as it is smaller and lacks an extension that overlaps with A12.2C in the Rpb9-like position (Fig. 2A). In Pol II, Rpb9C also binds the ED1, although in a different manner consistent with the structure of the Pol II ED1 (Fig. 2B). Therefore, Pol I and Pol II ED1 are specifically tailored to bind A12.2C and Rpb9C, respectively. Interestingly, a similar situation is observed in Pol III, where the ED2 is more structurally conserved than the ED1 (Fig. 2C). The Pol III ED1, as in Pol II, has an extension in the region where A12.2C binds, but the Rpb9C would also clash with an N-terminal helix from subunit C53. Accordingly, Pol III C11C (equivalent to Pol I A12.2C and Pol II Rbp9C) adopts a position near the jaw that differs from the position of both A12.2C and Rpb9C (Hoffmann et al., 2015) (Fig. 2C).

**Figure 2.**
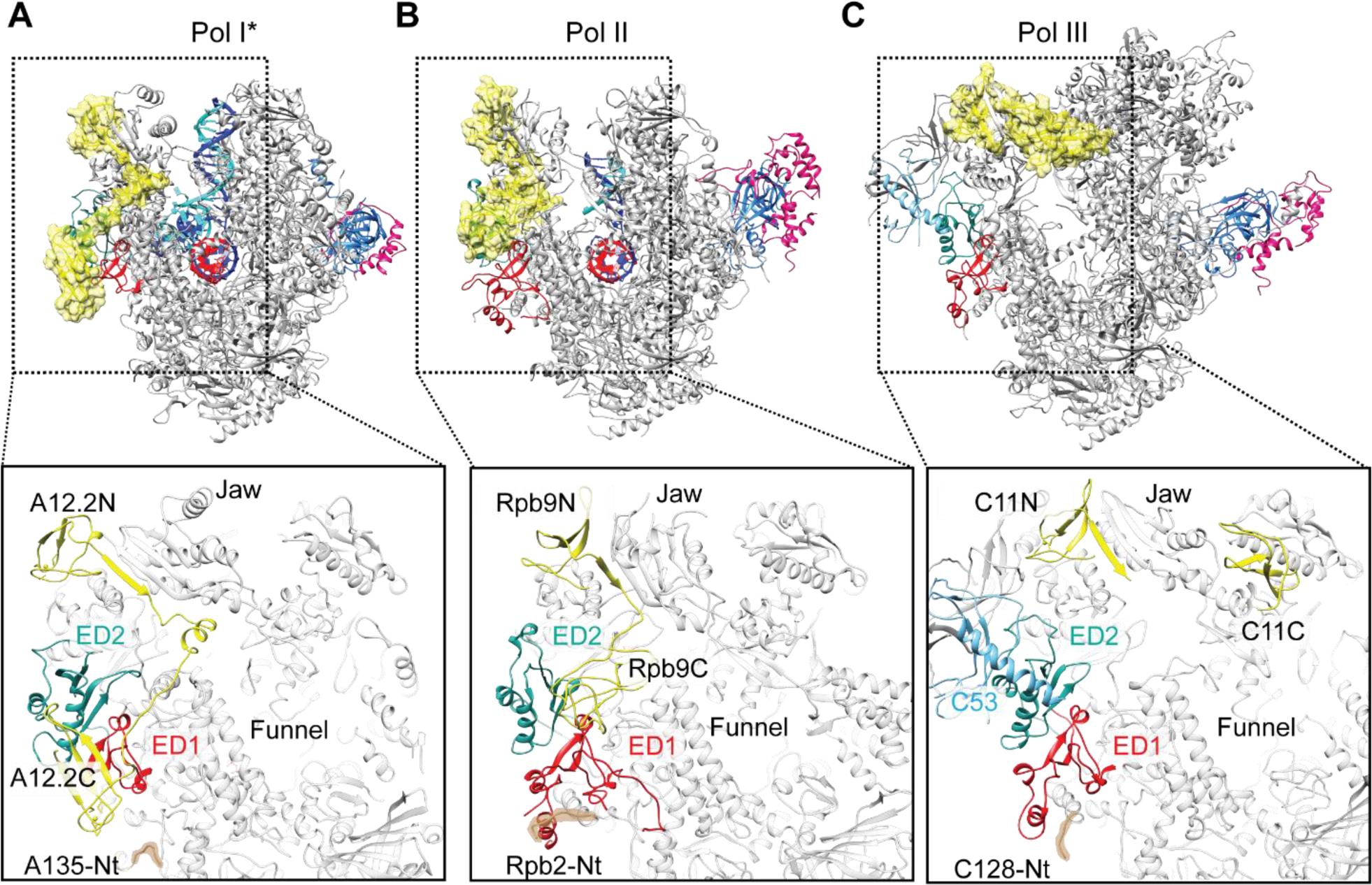
Comparison of the positions of the C-terminal domains of Pol I A12.2, Pol II Rbp9 and Pol III C11. The position of A12.2 (A), Rpb9 (B) or C11 (C) are shown in yellow for Pol I*, Pol II and Pol III, respectively. While the ED2 is structurally more conserved (light sea green color), the ED1 in Pol II and Pol III are larger than the Pol I ED1 (red). The structure of the ED1 determines the binding mode of Pol I A12.2C and Pol II Rpb9C, while in Pol III the presence of C53 induces a different binding site for C11C far from the ED. The position of the N-terminal tail of the second largest subunit is also indicated for each polymerase.

Further analysis of this region shows that the N-terminal tail of the Pol II second largest subunit (Rpb2), that forms part of the Rpb2 ED1 domain, is positioned similarly to the N-terminal region of C128, the corresponding Pol III subunit. The Pol I A135-Nt, however, is retracted and interacts with the HB domain as described previously, while in the presence of the heterodimer it is remains distant from the A135 EDs but moves away from the HB. This positioning allows binding of A12.2C to the ED1 and also controls the accessibility of the HB to the A34.5-Ct.

In conclusion, the A12.2C can only bind to the Rpb9-like position because of the distinct structure of the A135 ED1 and the different positioning of the A135-Nt. Heterodimer release and binding of A12.2C to the ED1 induce a conformational change in Pol I, which adopts a conformation resembling Pol II.

### NTP selection and discrimination are conserved among RNA polymerases

To better resolve smaller differences in the Pol I active site, particles from both EC reconstructions were pooled, and classification was restricted to the core enzyme and the DNA-RNA hybrid using a soft mask and higher weight on the data (Scheres, 2016) (Fig. 1-Figure Supplement 1, Material and Methods). This strategy allowed us to better resolve part of the partially “closed” trigger loop (TL), the path of the non-template strand in the transcription bubble (including the interactions on the downstream edge of the bubble), and the position and interactions of GMPCPP in the active site.

The NTP substrate is positioned in the “A” site, as previously seen in Pol II (Cheung, Sainsbury, & Cramer, 2011; Wang et al., 2006; Westover, Bushnell, & Kornberg, 2004) and bacterial RNA polymerase (bcPol)(Vassylyev et al., 2007) (Fig. 3A). Accordingly, the phosphate moiety is bound by two invariant arginine residues (A135 R714 and R957). In addition, two invariant residues N625 and R591, which are involved in NTP/dNTP discrimination, come close to the 3’- and 2’-OH group, respectively. While the corresponding residue to N625 in Pol II (Rpb1 N479) has been shown to interact with either the 3’-OH (Wang et al., 2006) or the 2’-OH (Cheung et al., 2011) group, the invariant R591 (Rpb1 R446) interacts with the 2’-OH group of the ribose in all Pol II structures. In addition, the NTP is maintained in the correct position by L1202 from the TL, which interacts with the guanosine base. Only up to this residue, weak density can be observed, while the “tip” loop (residues 1203-1212) is unresolved.

**Figure 3.**
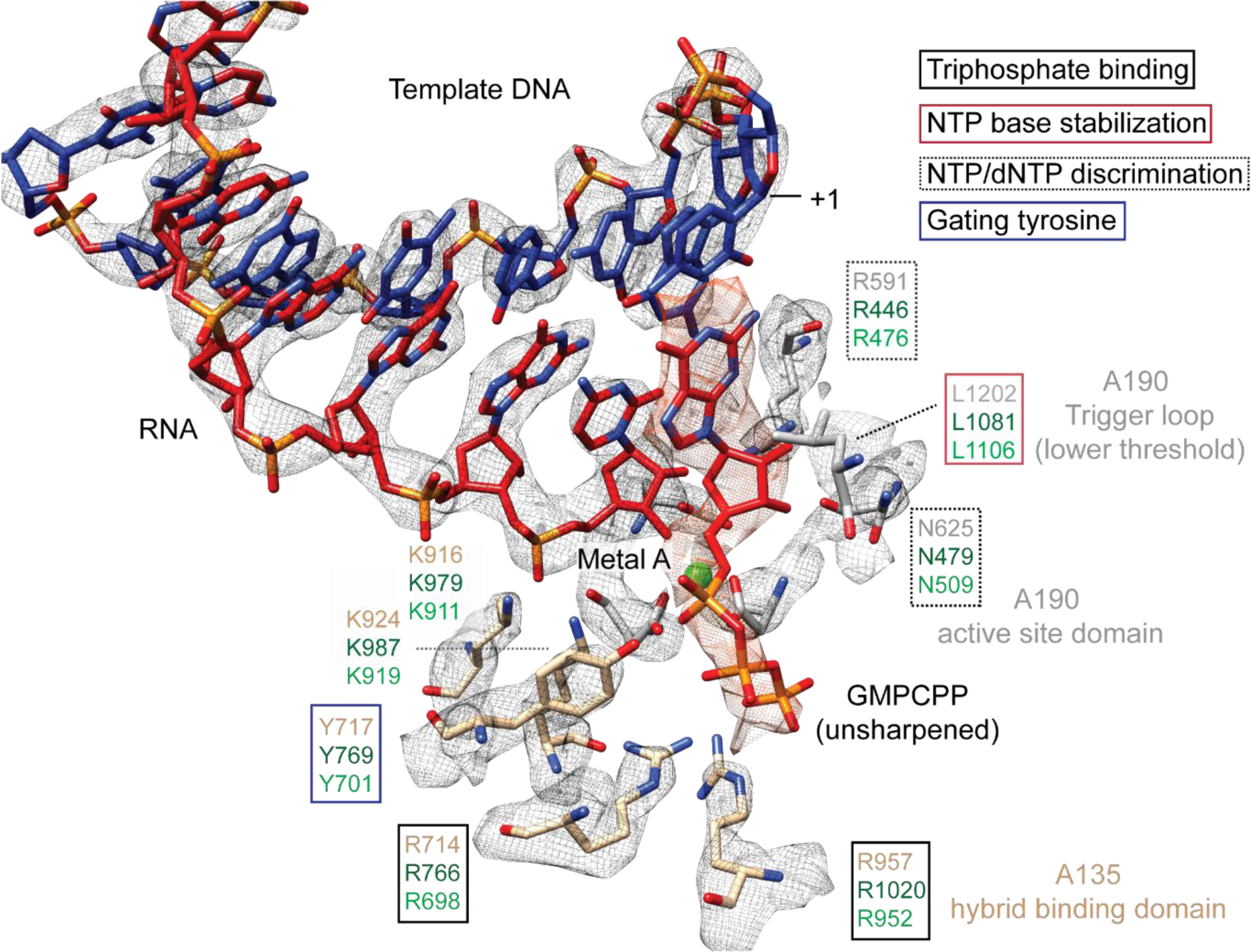
Density for the DNA-RNA hybrid, GMPCPP and its interactions with active site residues. GMPCPP is bound by conserved, identical residues in Pol I and Pol II. These include two arginines that interact with the phosphate (R714 and R957), a leucine from the trigger loop that stacks against the DNA base (L1202), and R591 and N625 which recognize the 2’- and 3’-OH groups, respectively. The “gating tyrosine” (Y717), involved in RNA positioning during backtracking(Cheung & Cramer, 2011), and K916 and K924, which bind the 3’-end of the RNA are also indicated. Residues are shown in grey (A190) or tan (A135) for Pol I, while those in Pol II in dark green, and in Pol III in light green. Density for the DNA-RNA hybrid is from the sharpened, Pol I (core) EC (+GMPCPP) reconstruction, while the GMPCPP is from the same reconstruction but from the unsharpened/unmasked map. Density for L1202 is shown at a lower threshold.

### Transcription bubble stabilization in Pol I

In the previous Pol I EC structures, the downstream DNA and the DNA-RNA hybrid adopted a conserved position compared to Pol II and Pol III (Neyer et al., 2016; Tafur et al., 2016). Additionally, the upstream DNA duplex was bound similar to the bovine Pol II (Bernecky, Herzog, Baumeister, Plitzko, & Cramer, 2016), but not to the available yeast Pol II structure (Barnes et al., 2015). In contrast to our previous data set^9^, the upstream DNA duplex is more flexible, but density could be improved by focused classification of the pooled Pol I* and Pol I ECs (Fig. 1-Figure Supplement 1). Two main classes were obtained that differed in the conformation of the rudder. One of them showed strong upstream DNA duplex density as well as density for the single-stranded non-template region (ssNT). While the lower resolution in this area precluded DNA sequence assignment and model building, continuous density from both ends of the transcription bubble delineate the path of the ssNT in the cleft (Fig. 1B, Fig. 4A). As suggested previously (Tafur et al., 2016), the ssNT strand follows a different path compared to the available structure of yeast Pol II (Barnes et al., 2015).

**Figure 4.**
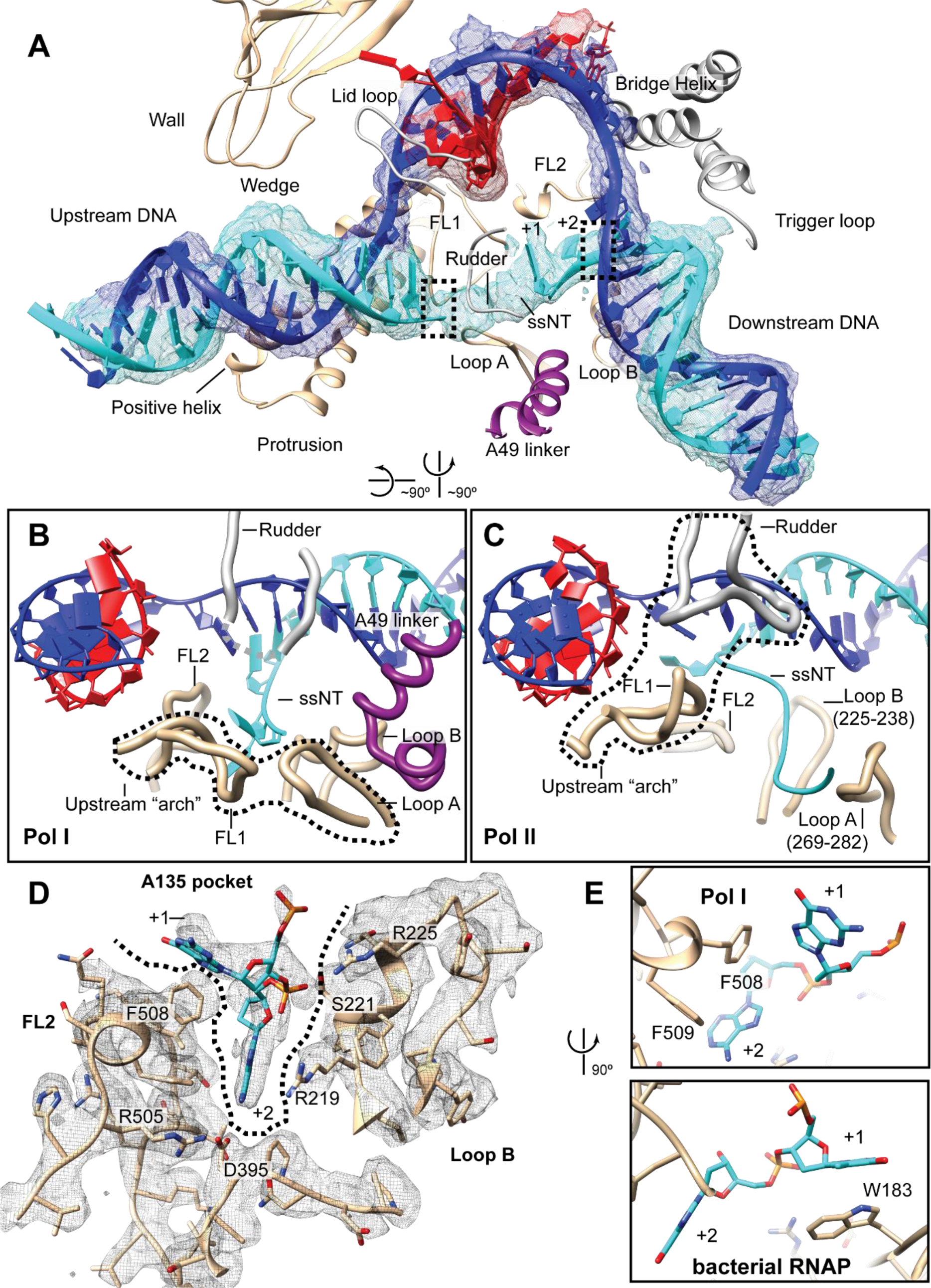
Interactions and stabilization of the transcription bubble in Pol I. **A.** Density for the full DNA/RNA transcription scaffold from the focused classification class is shown, next to the Pol I elements involved in binding the nucleic acids. The dotted boxes indicate the upstream and downstream boundaries of the transcription bubble. **B.** In Pol I, the “upstream arch” is composed of the open fork loop 1 (FL1) and loop A, while the rudder and the A49 linker helix only allow the passage of the single stranded non-template (ssNT) in a narrow cavity in between these elements. **C.** The upstream boundary of the transcription bubble in Pol II is determined by the rudder and closed FL1, which form an “upstream arch”. The corresponding regions to loop A and loop B are also indicated with numbering of the Rpb2 subunit. **D.** In the downstream edge of the transcription bubble, the +2 base of the NT strand is flipped into a pocket formed by FL2 and loop B (“A135 pocket”). These elements interact with the nucleotide through R219, R225 and the conserved D395. **E.** These interactions also position the +1 base next to F508 from FL2 (top), resembling the interaction of the +1 base with βW183 in bacterial Pol (bottom).

Although the ssNT density is clear, it is too short to accommodate all nucleotides between the transcription bubble boundaries. Additionally, the density becomes bulky near the rudder, suggesting that the ssNT strand wraps around the rudder or that it is in a strained conformation (Fig. 4-Figure Supplement 1). Altogether, these results suggest that the ssNT strand in the transcription bubble is dynamic and interacts intimately with the rudder. As the mismatch of 11 nucleotides was artificially induced into the transcription bubble, it is possible that the naturally occurring transcription bubble in active Pol I is shorter, although it is able to accommodate larger mismatches as other RNA polymerases (Barnes et al., 2015; Pal, Ponticelli, & Luse, 2005).

While the A135 protrusion positive helix and the wedge interact and stabilize the upstream DNA duplex, the rudder physically blocks the direction of the DNA duplex, defining the upstream boundary of the transcription bubble (Fig. 4A). The fork loop 1 (FL1) is tilted down towards the A135 lobe and forms an “upstream arch” with loop A, which physically restricts the passage of the ssNT strand (Fig. 4B). In contrast, in Pol II, the “upstream arch” is composed of the rudder and the FL1(Barnes et al., 2015) and is differently configured (Fig. 4C). The corresponding region to loop A (Rpb2 residues 269-282) is retracted, allowing the positioning of the ssNT strand in between this region and the “upstream arch”. In Pol I, the configuration of the “upstream arch” only allows the positioning of the ssNT strand in a space in between the FL1, loop A and the rudder. In the context of the 14-subunit Pol I, further restriction of the allowable position of the ssNT strand is enforced by the A49 linker helix that spans the cleft (Fig. 4A, B).

In Pol II, the downstream edge of the bubble is determined by the FL2, which similarly to the FL1, can alternate between “open” and “closed” states (Barnes et al., 2015). When the FL2 is in an “open” conformation, the +2 base of the ssNT strand is flipped (Cheung & Cramer,2010) (Fig. 4-Figure Supplement 2). In the yeast Pol II structure with a complete transcription bubble, however, the FL2 adopts a “closed” conformation, and shifts the position of the +2 base in the ssNT strand(Barnes et al., 2015) (Fig. 4-Figure Supplement 2). In bcPol, nucleotide +2 from the ssNT strand is flipped and inserted into a pocket formed by residues from the β subunit (“β pocket”) (Zhang et al., 2012). Interestingly, in the Pol I EC, the +2 nucleotide is also flipped into a narrow pocket (“A135 pocket”) formed by loop B and the FL2, which is in an “open” conformation (Fig. 4D). While residues from the FL2 face the flipped base, loop B exposes positively charged residues to the nucleobase accommodated in the pocket. In particular, R219 and R225 appear to stabilize the flipped +2 base, promoting the stacking of the +1 base with F508 (from FL2) (Fig.4E, top). The interaction of the +1 base with F508 is reminiscent of the interaction of the same base with W183 of subunit β of bcPol (Zhang et al., 2012) (Fig.4E, bottom). In both bcPol and Pol I, the interactions with the +1 and +2 nucleotides appear to stabilize and direct the ssNT strand in the cleft. Additionally, the highly conserved D395 also interacts with the +2 base as in bcPol βD446) (Vvedenskaya et al., 2014) and Pol II (Rpb2 D399) (Cheung & Cramer, 2011), and probably also in Pol III (C128 D370). F508 is not conserved in the FL2 of Pol II but is present in the Pol III FL2 (C128 F477). However, neither Pol II nor Pol III can form the equivalent interactions as in the A135 pocket because their corresponding loop B is differently positioned and far from the +1 and +2 bases.

Classification of the pooled consensus ECs (Fig. 1-Figure Supplement 1) by the conformation of the core revealed a continuum of states in which the +2 base was either not flipped (when the cleft was slightly more open) or flipped (when the cleft was completely closed) (Fig. 4-Figure Supplement 3). Interestingly, flipping of the +2 base was only observed when the jaw and clamp domains rotated relative to each other before the complete closure of the cleft and coinciding with DNA-RNA hybrid stabilization, stable binding of the NTP and partial TL closing (transition from state 3 to 4 in Fig. 4-Figure Supplement 3; Video 1). Thus, +2 base flipping only occurs when the complex is correctly positioned for catalysis. The variability in the position of this base could also explain why in previous Pol I EC structures, in addition to the lower resolution and lower number of particles, this base was modelled unflipped (Tafur et al., 2016) or not modelled (Neyer et al., 2016). Strikingly, comparison of these reconstructions shows that formation of the final EC requires the concerted, sequential movement of different domains (Video 2). Upon DNA-RNA binding, the protrusion and wall domains move towards the clamp, followed by the jaw/clamp movement that promotes +2 base flipping and finally closing of the cleft by movement of the complete modules 1 and 2.

## Discussion

After the crystal structure of the Pol I dimer was published (Engel et al., 2013; Fernandez-Tornero et al., 2013), the active enzyme in its initiating (Engel et al., 2017; Han et al., 2017; Sadian et al., 2017) and elongating (Neyer et al., 2016; Tafur et al., 2016) forms have been solved by cryo-EM. While the general arrangement of subunits is similar in all reconstructions, one of the main differences between them is the position of A12.2C. In the active form of the enzyme, A12.2C is excluded from the active site, but its alternative position could not be determined. Here, we show that A12.2C can alternate between TFIIS-like and Rpb9-like positions, partially depending on the presence of the heterodimer. In the TFIIS-like position, A12.2C is in the cleft and occludes the active site, thus being incompatible with NTP incorporation. When the cleft is closed (and thereby clashes with A12.2C in the TFIIS-like position), A12.2C is excluded from the active site and can bind to a distinct site near the A135 EDs. Binding to the ED1 requires the release of the heterodimer in the complex, as both the A12.2C and the A34.5-Ct bind to the ED1 using overlapping binding sites. Release of the heterodimer is also promoted by the movement of the A135-Nt towards the HB, which blocks the interaction of the distal part of the A34.5-Ct with this domain. These results suggest a mechanism by which the surface of A135 (in particular, the ED1) plays a pivotal role in specific factor exchange in Pol I. It appears that in both Pol I and Pol II, the configuration of the ED1 is such that it can only accommodate their respective Rpb9-like C-terminal domain, although in different binding modes. In contrast, in Pol III, the presence of C53 prevents the binding of C11C to the C128 ED1, favoring its positioning near the jaw. In addition, while in both Pol II and Pol III the N-terminal region of the second biggest subunit is part of the ED1, in Pol I this flexible tail is located distant from the ED1 and functions as a switch which alternates its position depending on the presence or absence of the A34.5-Ct. In combination with the binding of A12.2C to the ED1, the interaction of the A135-Nt with the HB “seals” two of the three main binding interfaces of A34.5-Ct with Pol I (the third being the anchoring of the very end of A34.5 to a cavity formed by A135, AC40 and Rpb10). Therefore, in Pol I*, the heterodimer is excluded from the enzyme and Pol I adopts a Pol II-like conformation.

Our results show that Pol I can adopt an active conformation (with a stable DNA-RNA hybrid, +2 base flipped and the NTP in the active site) in the absence of the heterodimer. With our data, however, it is not possible to discern whether the heterodimer is excluded before or after formation of this complex. Based on the extensive interaction surface between the A34.5-Ct and A135, it is likely that the heterodimer dissociates after the formation of a stable EC, which allows A12.2C binding to the ED1. Recent genetic studies have suggested that A12.2 may be also involved in modulation of the movement of the jaw/lobe interface especially in the absence of A49, as the linker and tWH domains appear to stabilize the closed conformation of Pol I when bound to DNA (Darrière T, 2018). As the A12.2C binds to the A135 ED1, which sits next to the A135 lobe, the A12.2C might restrict the movement of the lobe. Thus, while A12.2N regulates the flexibility of the jaw, A12.2C can additionally regulate the movement of the lobe. Together, both A12.2 domains could therefore regulate cleft opening/closing of Pol I upon DNA binding. Restriction of movement of the A135 lobe by the A12.2C might be important to maintain the closed state in the absence of A49, as in Pol I*. A requirement for the movement of the jaw/lobe during Pol I elongation is further supported by the fact that +2 base flipping (and subsequent transcription bubble stabilization) is established only when the jaw and clamp domains move relative to each other.

*In vivo*, heterodimer association to Pol I might offer an additional layer of regulation of rDNA transcription (Figure 5). The proportion of initiation-competent Pol I molecules in the cell has been proposed to represent those Pol I particles bound to initiation factor Rrn3 (Milkereit & Tschochner, 1998). Conversely, the number of Pol I* particles in the cell could represent a population of actively transcribing DNA-bound Pol I, but also a pool of readily active Pol I that can initiate transcription upon heterodimer binding (in contrast to Pol I dimers, which appear to be a storage form of the enzyme (Torreira et al., 2017)). The number of initiation-competent Pol I molecules could be thus regulated not only by Pol I homo-dimerization and association with Rrn3, but also by changes in the heterodimer concentration in the nucleolus, thereby controlling the ratio of Pol I to Pol I*. Nutrient-dependent regulation of nucleolar localization of the mammalian A49-A34.5 homolog PAF53-PAF49 has been observed (Penrod et al., 2015). PAF49 (A34.5 counterpart) accumulates in the nucleolus in growing cells but disperses to the nucleoplasm upon serum starvation (Yamamoto et al., 2004). In yeast, A34.5 is maintained in the nucleolus by its association with A49 (but also contains a nucleolar localization signal in its C-terminal region), and A49 is required for the high loading rate of Pol I onto rDNA (Albert et al., 2011). Human PAF53-PAF49 can substitute the A49-A34.5 heterodimer *in vivo* (Albert et al., 2011), suggesting a conserved function (and possibly regulation). Regulation of heterodimer binding to Pol I might also explain why promoter association of Pol I-Rrn3 complexes is low upon nutrient starvation even when the concentration of such complexes is relatively high (Torreira et al., 2017); the levels of the heterodimer might further regulate Pol I initiation rates.

Interestingly, the release of the heterodimer from the enzyme would also allow the binding of elongation factors to Pol I. Pol I has been shown to bind Spt5 directly (Viktorovskaya, Appling, & Schneider, 2011) and its activity is affected by Spt4/5 *in vivo* (Anderson et al., 2011). In the Pol I EC, canonical binding of Spt4/5 is precluded by the A49 tWH (Tafur et al., 2016), as it occupies a position equivalent to the KOW1-L1 domain of Spt5, and by the A49 linker helix spanning the cleft, which clashes slightly with the N-terminal region of Spt5 (Bernecky, Plitzko, & Cramer, 2017; Ehara et al., 2017). Interestingly, Spt5 interacts physically and genetically with A49, suggesting a functional interplay between these proteins (Viktorovskaya et al., 2011). Paf1C, another elongation factor, has also been shown to stimulate Pol I transcription *in vivo* and *in vitro* (Zhang, Sikes, Beyer, & Schneider, 2009; Zhang, Smith, Renfrow, & Schneider, 2010). Paf1C binds to Pol II on the outer surface of subunit Rpb2 (Pol II counterpart of A135), including the Rpb2 ED2 and lobe (Xu et al., 2017). In this position, it clashes and competes with TFIIF for Pol II binding (Xu et al., 2017). Heterodimer dissociation from Pol I frees the binding site for both Spt4/5 and Paf1C in a mechanism that could be akin to the transition from initiation to elongation in Pol II: while TFIIE (A49 tWH) blocks the Spt4/5 binding site, TFIIF (A49-A34.5 dimerization domain) occupies the binding site of part of Paf1C (Xu et al., 2017) (Fig. 5-Figure Supplement 1). Thus, binding of elongation factors is mutually exclusive with the presence of initiation factors. Therefore, in Pol I, factor exchange during the transition from initiation to elongation could be accommodated more readily just by the release of the heterodimer and switching to the Pol II-like Pol I* form. A similar allosteric transition during promoter escape in Pol I mediated by the heterodimer, Spt5 and the stalk has been previously proposed (Beckouet, Mariotte-Labarre, Peyroche, Nogi, & Thuriaux, 2011).

**Figure 5.**
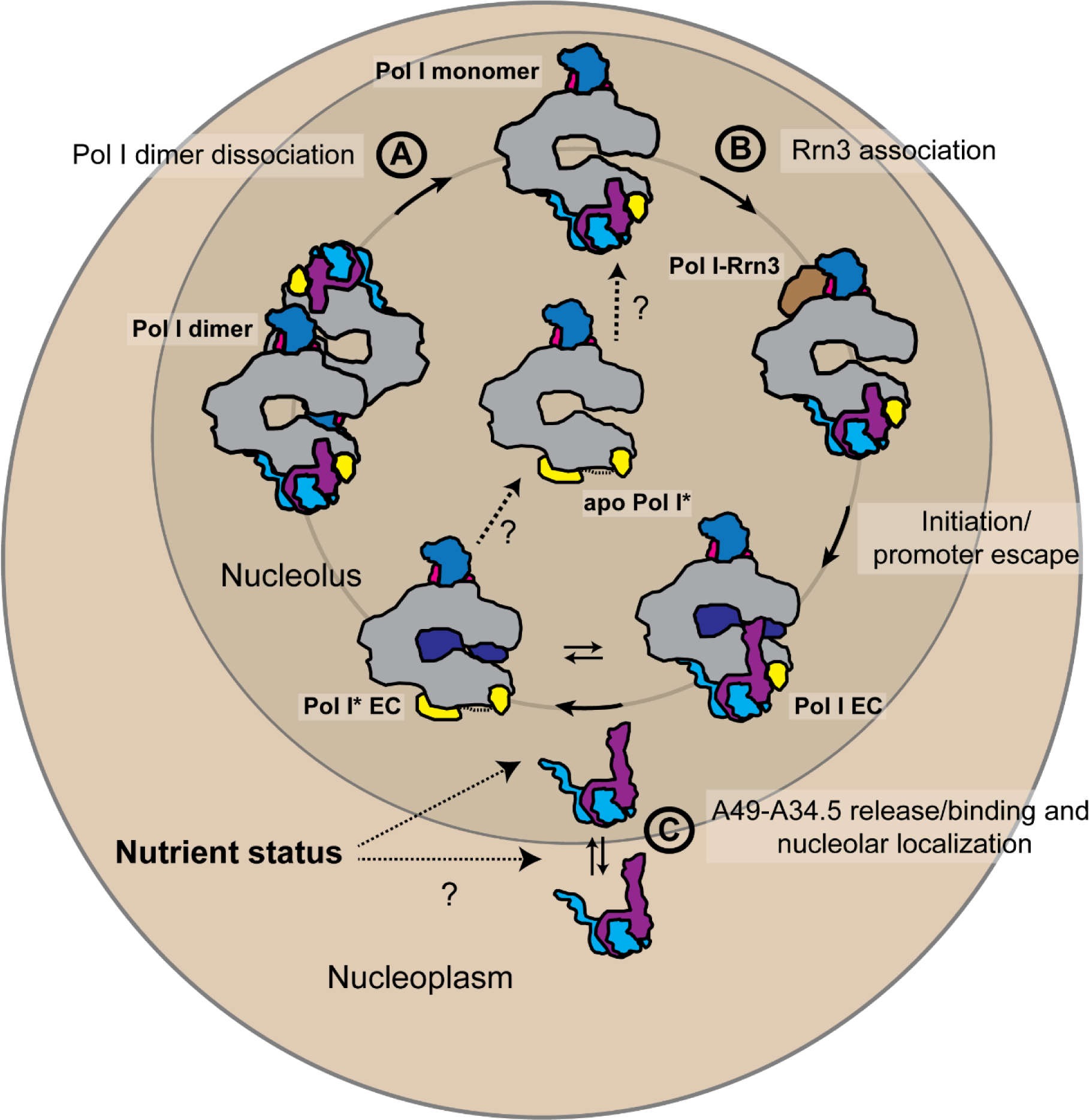
Schematic representation of the possible physiological role of the A49-A34.5 heterodimer in the regulation Pol I activity. The pool of initiation-competent Pol I particles is controlled by Pol I homo-dimerization (A) and binding of Rrn3 to monomeric Pol I (B). After transcription initiation and promoter escape, during elongation, Pol I can alternate between Pol I and Pol I* conformations. Release of the A49-A34.5 heterodimer would allow the recruitment of elongation factors (C). After dissociating from DNA, Pol I* could bind to the A49-A34.5 heterodimer to replenish the pool of initiation-competent Pol I monomers. The concentration of A49-A34.5 heterodimer in the nucleolus might be also regulated by the nutrient status of the cell as in the mammalian system. Regulated localization of the A49-A34.5 heterodimer would serve to alter the ratio of Pol I to Pol I* in the nucleolus, thereby controlling the initiation rates on the rDNA.

Given the conservation of residues in the active site among RNA polymerases, similar binding of the NTP as previously observed in Pol II (Wang et al., 2006; Westover et al., 2004) and bcPol (Vassylyev et al., 2007) was expected. Accordingly, in our structures, GMPCPP is positioned in the “A” site by two invariant arginines (A135 R714 and R957), which interact with the phosphate moiety, and by TL residue L1202 which stacks with the base. These interactions stabilize the base pairing of the incoming NTP with the +1 nucleotide, promoting NMP incorporation. Stabilization of the NTP in the Pol I active site by L1202 (Huang et al., 2010) suggests that, as in Pol II, the TL effectively “seals” the active site and acts as a “positional catalyst” (Mishanina, Palo, Nayak, Mooney, & Landick, 2017). In addition, both conserved N625 and R591 are positioned close to the 2’ and 3’-OH groups, in agreement with their role on NTP/dNTP discrimination. Quantum-chemical analysis of NTP/dNTP discrimination in Pol II has shown that R446 (equivalent to Pol I R591) is critical (Rossbach & Ochsenfeld, 2017), explaining why its mutation is lethal (Wang et al., 2006). Altogether, the same set of interactions observed between the NTP and residues from the active site in Pol II are also observed in Pol I, suggesting a universal mechanism of catalysis. Interestingly, transient-state kinetic analysis of yeast Pol I transcription has revealed that upon NTP binding, two different Pol I populations are observed (Appling, Lucius, & Schneider, 2015). Additionally, A12.2 appears to play a role during nucleotide incorporation which differs from its nucleolytic activity (Appling, Schneider, & Lucius, 2017). Therefore, these experiments are in agreement with our observations under cryo-EM conditions and suggest that upon NTP binding, Pol I can conformationally switch to Pol I*, which in turn affects the catalytic properties of the enzyme.

Our structures also reveal how the active, elongating complex stabilizes the transcription bubble. Of particular interest is the interaction of the +1 and +2 bases with residues from the FL2 and loop B. In a similar manner to what has been observed in bcPol, the +2 base plays an important role in stabilization of the transcription bubble at the downstream edge (Zhang et al., 2012). In bcPol, the +2 base is part of the core recognition element (CRE) and recognition is base-specific towards a guanine (+2G) (Zhang et al., 2012). Interactions with the CRE and +2G are important for transcription start site selection (Vvedenskaya et al., 2016) and stability of the open complex (Zhang et al., 2012), and these interactions appear to also occur during elongation (Vvedenskaya et al., 2014). Mutation of βD446, which recognizes the guanine, affects positively or negatively transcriptional pausing, depending on the nature of the pause (Petushkov, Pupov, Bass, & Kulbachinskiy, 2015; Vvedenskaya et al., 2014). Conservation of D395 in yeast and bacterial Pols suggests that +2 base flipping is a conserved mechanism to stabilize the downstream, leading edge of the transcription bubble. In Pol I, stabilization of the active EC might also be dependent on the extensive interactions of residues from the A135 pocket with the +2 base, which include, in addition to D395, R219 and R225 from loop B.

The interactions of Pol I with the +1 and +2 base appear to direct the non-template strand between the A49 linker helix, the rudder and the “upstream arch” composed of FL1 and loop A. Base stacking of F508 with the +1 nucleotide might be a conserved feature adapted from bcPol (which uses βW183 to interact with the +1) to direct the ssNT in the cleft. Mutational analysis of W183 has revealed that it plays a role during the initial steps of RNA synthesis and during translocation in the *E.coli* σ^54^ system, preventing the formation of short abortive transcripts (Wiesler, Weinzierl, & Buck, 2013). A similar scenario in Pol I, whereby F508 promotes the synthesis of productive versus abortive transcripts, might stimulate the high processivity needed due to the high initiation rates on the rDNA. It is tempting to speculate that equivalent interactions as the CRE-Pol in bacteria also occur during Pol I initiation as the nature of the +1 (pyrimidine) and +2 (purine) is conserved in the yeast rDNA sequence and we see similar interactions of Pol I with both bases. A role of these residues during transcription initiation awaits further structures of Pol I initiation complexes.

Given the recent characterization of the pre-initiation machineries in Pol I (Engel et al., 2017; Han et al., 2017; Sadian et al., 2017), Pol II (He et al., 2016; Plaschka et al., 2016) and Pol III (Abascal-Palacios et al., 2018; Vorlander et al., 2018), it appears that while the assembly of the pre-initiation complex is more divergent and specialized, Pol I displays features from both Pol II and bcPol during elongation, arguing in favor of a universal catalytic mechanism in RNA polymerases.

## Material and Methods

### Pol I EC-GMPCPP complex formation

Endogenous Pol I was purified from yeast cells as previously described (Moreno-Morcillo et al., 2014). Pol I was incubated with a 38 base pair transcription scaffold containing an 11 nucleotide mismatch bubble and a 20 nucleotide RNA as used previously for formation of the Pol I EC (Tafur et al., 2016). The complex was incubated for 1 hour at 4°C in 15 mM HEPES-NaOH (pH 7.5), 150 mM ammonium sulfate, 1 mM MgCl_2_, 1 mM GMPCPP (Jena Bioscience) and 10 mM DTT. The sample was diluted to ~0.1 mg/mL in the same buffer immediately before grid freezing.

### Cryo-EM sample preparation

2.5 μL of sample was deposited on a freshly glow-discharged cryo grid (R 2/1 + 2nm carbon, Quantifoil), incubated for 30 s, and blotted for 3 s (with a blotting force of “3”), at 100% humidity and 4°C in a Vitrobot Mark IV (FEI). Grids were stored in liquid nitrogen until data collection.

### Cryo-EM data collection

5,768 micrograph movies were collected on a FEI Titan Krios at 300 keV through a Gatan Quantum 967 LS energy filter using a 20 eV slit width in zero-loss mode. The movies were recorded on a Gatan K2 direct electron detector, at a nominal magnification of 135,000x corresponding to a pixel size of 1.04 Å in super resolution mode, using Serial EM. Movies were collected in 40 frames with defocus values from -0.75 to -2.5 μM, with a dose of 0.9775 e^−^ Å^−2^ s^−1^ per frame for 16 s.

### Cryo-EM data processing

Movies were aligned, motion-corrected and dose-fractionated using MotionCor2 (Zheng et al., 2017). Contrast transfer function (CTF) estimation was done using CTFFIND4 (Rohou & Grigorieff, 2015). All processing steps were performed in Relion 2.0 (Kimanius et al., 2016) unless otherwise indicated. Resolution estimates reported are those obtained after masking and B-factor sharpening (Relion post-processing). Data were divided in five batches to increase processing speed. For each batch, autopicking was followed by a 2D classification step (with data downsized 5 times) to remove contamination and damaged particles. Good classes were selected, re-extracted and un-binned, and refined against the Pol I EC (PDB: 5m5x) low pass filtered to 40 Å. Then, a 3D classification step was performed without alignment. For all batches the same procedure was followed, except for batch 5, in which 3D classification was performed with data downsized 5 times. Classes were selected based on the width of the cleft, the position of the clamp, and the DNA-RNA scaffold density, and grouped by similarity. Refinement of the pooled particles with closed cleft and strong DNA-RNA density revealed an extra density and streaky, weak density for the A49-A34.5 heterodimer. To resolve this region, a masked classification was performed. This yielded a class with high resolution in the extra density, allowing the unambiguous assignment of the A12.2 C-terminal domain (A12.2C). Based on these results, all other pooled classes were classified with a mask on this area. Particles were merged depending on whether they showed density for the A49-A34.5 heterodimer (Pol I) or the A12.2C without A49-A34.5 (Pol I*). During the process, additional bad particles were discarded by global 3D classification without a mask nor alignment. After refinement of all good particles for Pol I and Pol I*, additional classification steps were performed to increase the resolvability of the active site. For Pol I* particles, a 3D classification step with a mask on the core and DNA-RNA hybrid yielded a class (182,488 particles) with a better density for GMPCPP, which could be refined to 3.18 Å resolution. An apo form of Pol I* consisting of 73,660 particles was obtained during a global classification step of the initial subset with a closed cleft and strong DNA-RNA density, and was refined to 3.21 Å resolution. For the pooled Pol I particles, a global 3D classification step yielded a class with a closed clamp (EC) and a class bound to DNA-RNA with a slightly more open clamp. The latter was classified one more round, which gave a class in an EC conformation. Thiese particles weres merged with the EC particles from the previous 3D classification step, refined (consensus Pol I EC) and classified with a mask on the core, the full DNA-RNA scaffold and the linker helix of A49, which yielded a class with strong GMPCPP density (30,232 particles) that was refined to 3.42 Å resolution. As both Pol I EC and Pol I* EC reconstructions were very similar in the active site, EC particles were merged and classified using different masks. Masked classification based on the full DNA scaffold and rudder produced one class (34,475 particles) with improved density for the upstream DNA duplex and revealing the path of the single stranded non-template strand (ssNT), which was refined to 4.0 Å resolution (without post-processing). Classification based on the core and DNA-RNA scaffold revealed different states differing in the width of the cleft, base flipping at position +2, presence of the GMPCPP and conformation of the trigger loop (shown in Fig 4.-Figure Supplement 3). One of these classes (Pol I (core) EC +GMPCPP), which showed better density for GMPCPP, the +2 base and A190 L1202 was refined to 3.18 Å resolution (54,017 particles). Local resolution was estimated with Blocres (Cardone, Heymann, & Steven, 2013).

### Model building and refinement

Previous Pol I structures in its apo (PDB: 4c3i and 4c2m) and elongating (PDB: 5m5x) forms were used as starting models. The initial placement of GMPCPP in the active site was based on its position in a Pol II EC with bound GMPCPP (Wang et al., 2006) (PDB: 2e2j). Initially, the model for the Pol I (core) EC (+GMPCPP) was built in COOT (Emsley & Cowtan, 2004) and real-space refined in PHENIX (Adams et al., 2010). This model was then rigid body fitted in the Pol I* or Pol I EC (+ GMPCPP) maps in UCSF Chimera, further adjusted in COOT, and real-space refined again in PHENIX. For Pol I*, residues 66-125 from A12.2 were taken from the apo crystal structure (PDB: 4c3i), fitted to the density and manually adjusted. The A12.2 linker region was deleted afterwards. Agreement between maps and models was estimated in PHENIX. Model quality was assessed with Molprobity(Chen et al., 2010).

### Accession numbers

Models have been deposited in the PDB with codes: XXX (Pol I (core) EC +GMPCPP), XXX (Pol I EC +GMPCPP) and XXX (Pol I* EC +GMPCPP), while cryo-EM maps have been deposited with codes: XXX (Pol I (core) EC +GMPCPP), XXX (Pol I EC+GMPCPP), XXX (Pol I* EC+GMPCPP), XXX (Pol I (core) EC-upstream DNA focus), and XXX (apo Pol I*).

## Acknowledgements

Y.S, L.T, R.W and C.W.M acknowledge support by the ERC Advanced Grant (ERC-2013-AdG340964-POL1PIC). L.T acknowledges support by the EMBL International PhD program. We thank Jonas Hanske for discussion and comments on the manuscript.

## Author’s contributions

C.W.M initiated and supervised the project. L.T designed and carried out experiments, data processing and model building. Y.S aided in cryo-EM sample preparation. Y.S and F.W collected the cryo-EM data, with input from L.T. L.T analysed the data and built the models. R.W carried out yeast fermentation. L.T and C.W.M prepared the manuscript with input from all other authors.

---

**Figure 1-Figure Supplement 1.**
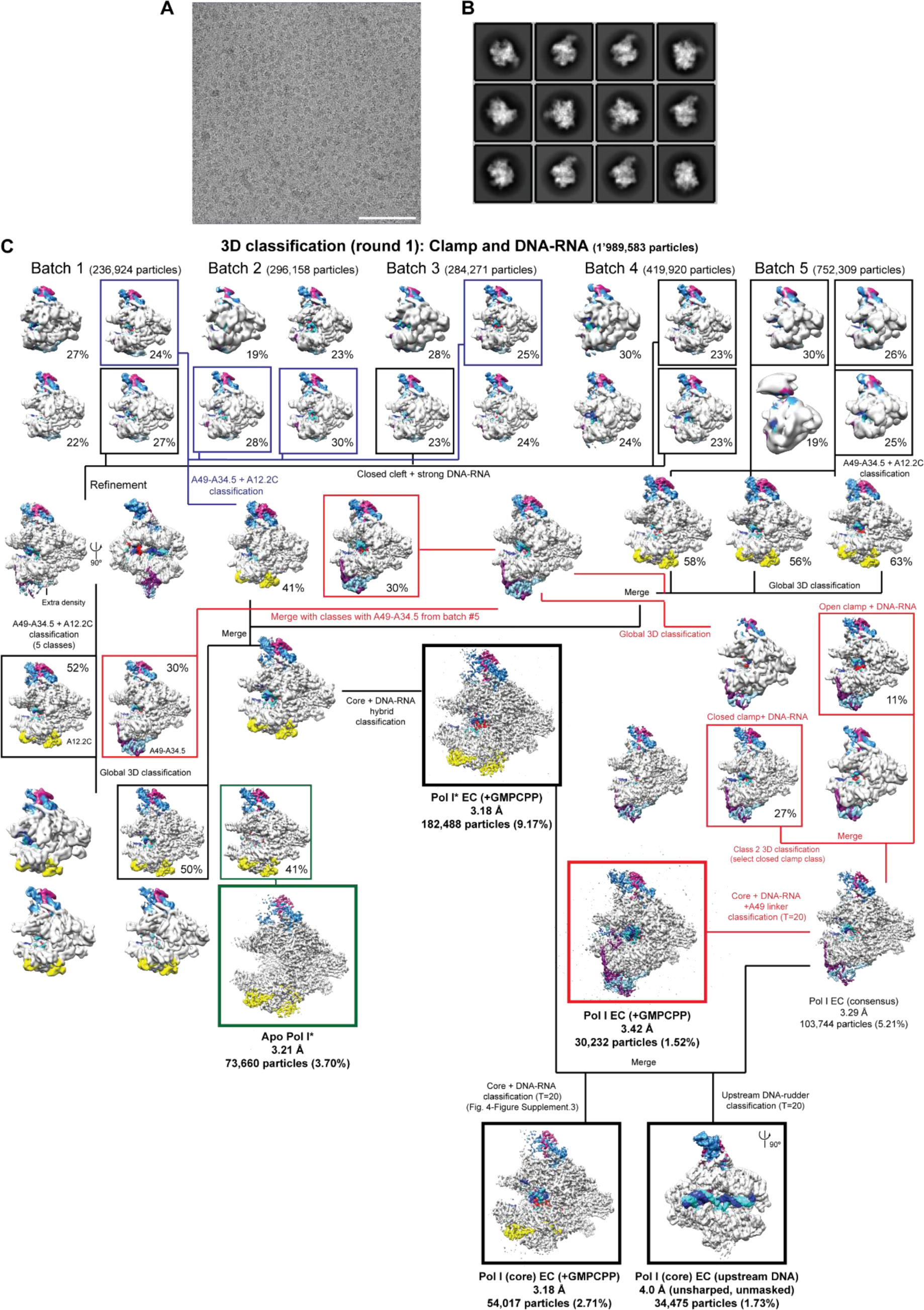
Cryo-EM data and processing. **A.** Representative micrograph. Scale bar = 100 nm. **B.** 2D class averages from the initial auto-picked particles. **C.** Processing pipeline. The resolution, number of particles and percentage of particles with respect to the initial number of particles after 2D classification is shown below each class.

**Figure 1-Figure Supplement 2.**
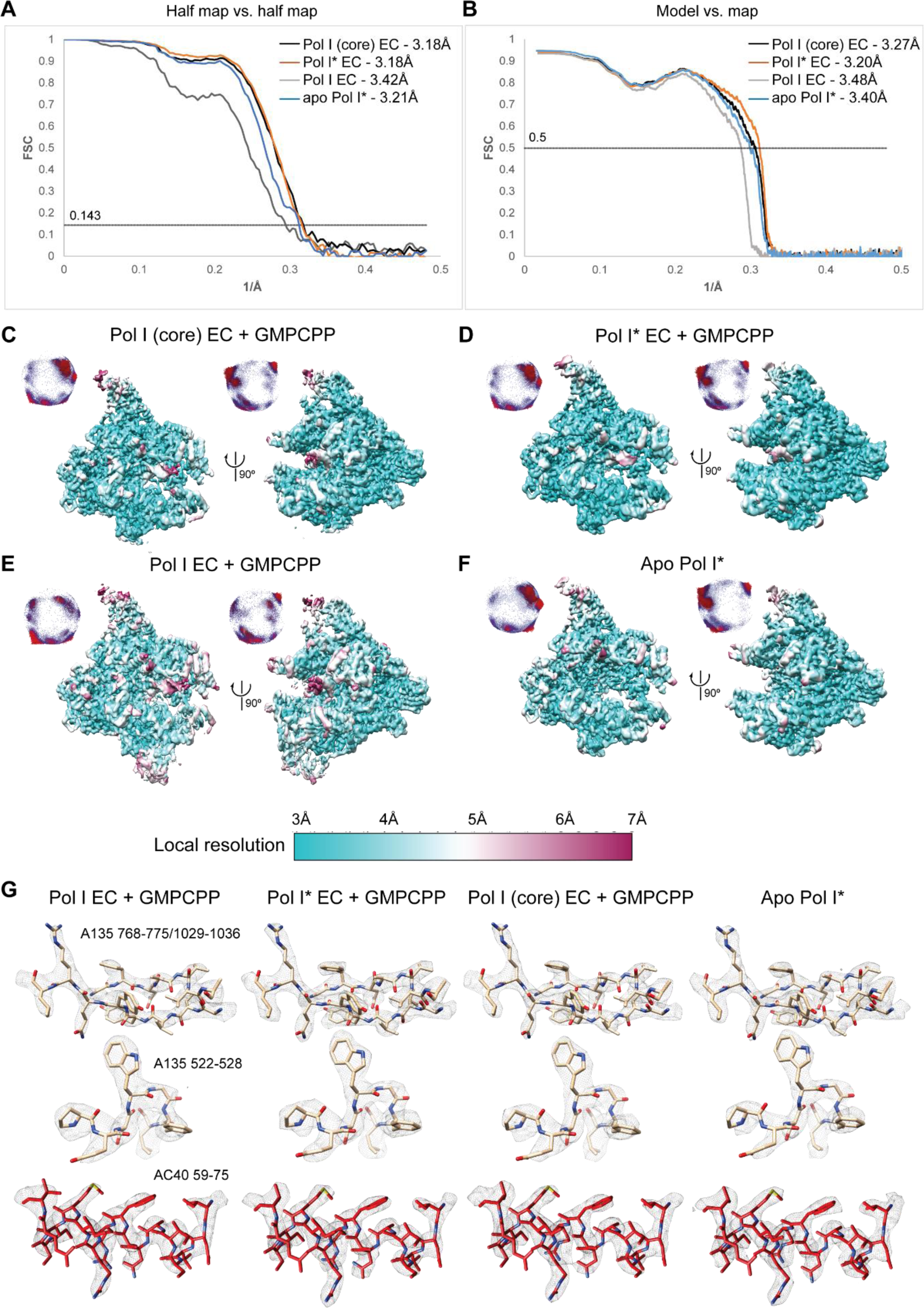
Average and local resolution estimates for the reconstructions. **A.** Fourier-shell correlation (FSC) curves for the reconstructions. **B**. FSC curves for the models versus experimental maps. **C-F.** Local resolution estimates for all the reconstructions. **G**. Representative densities.

**Figure 1-Figure Supplement 3.**
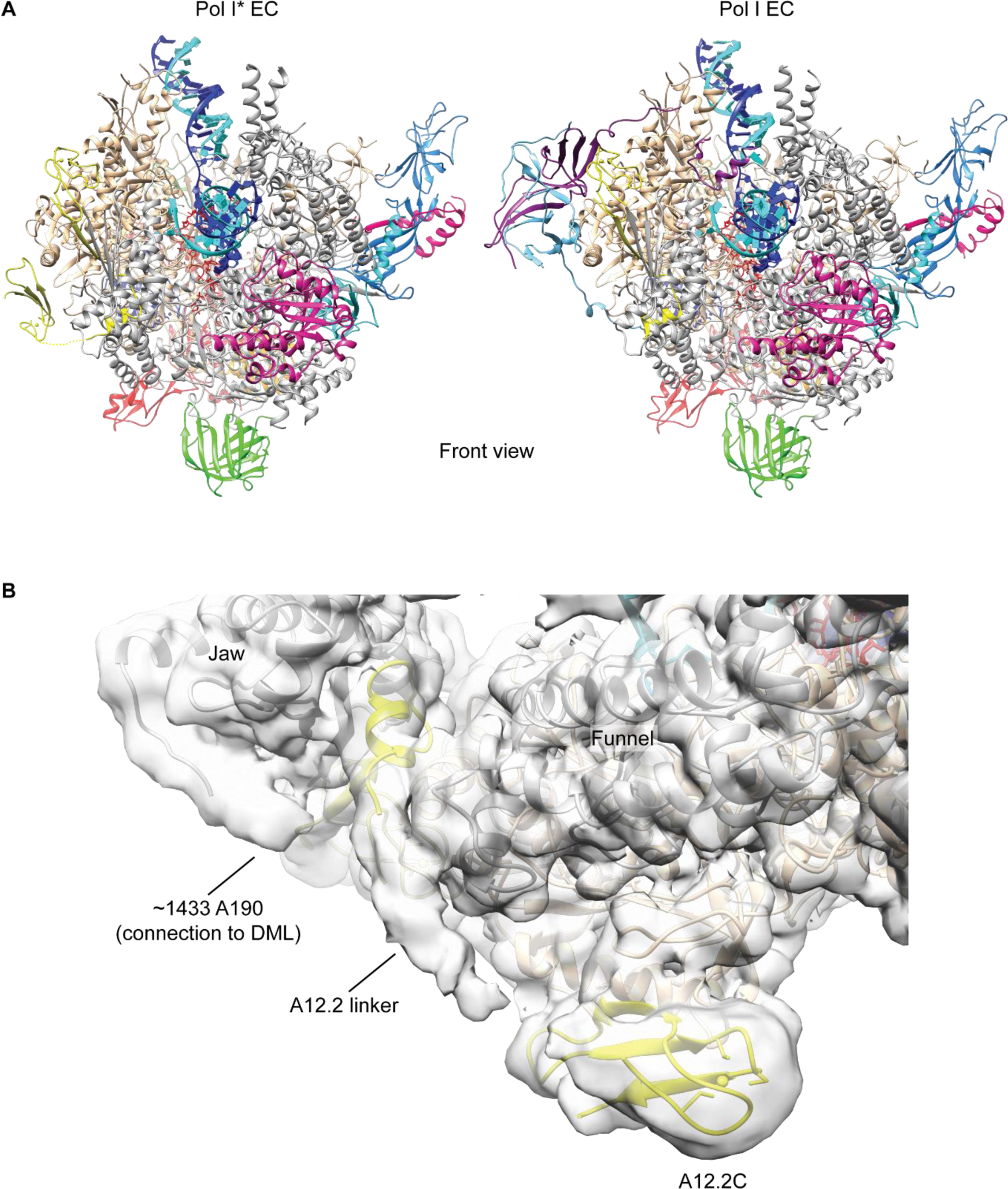
Pol I* and Pol I EC models. **A.** Front view of the Pol I* EC (left) and Pol I EC (right) models. **B**. Density for the A12.2 linker that connects the N-terminal and C-terminal Zn ribbons is visible at low threshold in the unsharpened map (Pol I* EC map is shown). Extra density from the jaw in the region that connects to the DNA-mimicking loop (DML) is also indicated.

**Figure 1-Figure Supplement 4.**
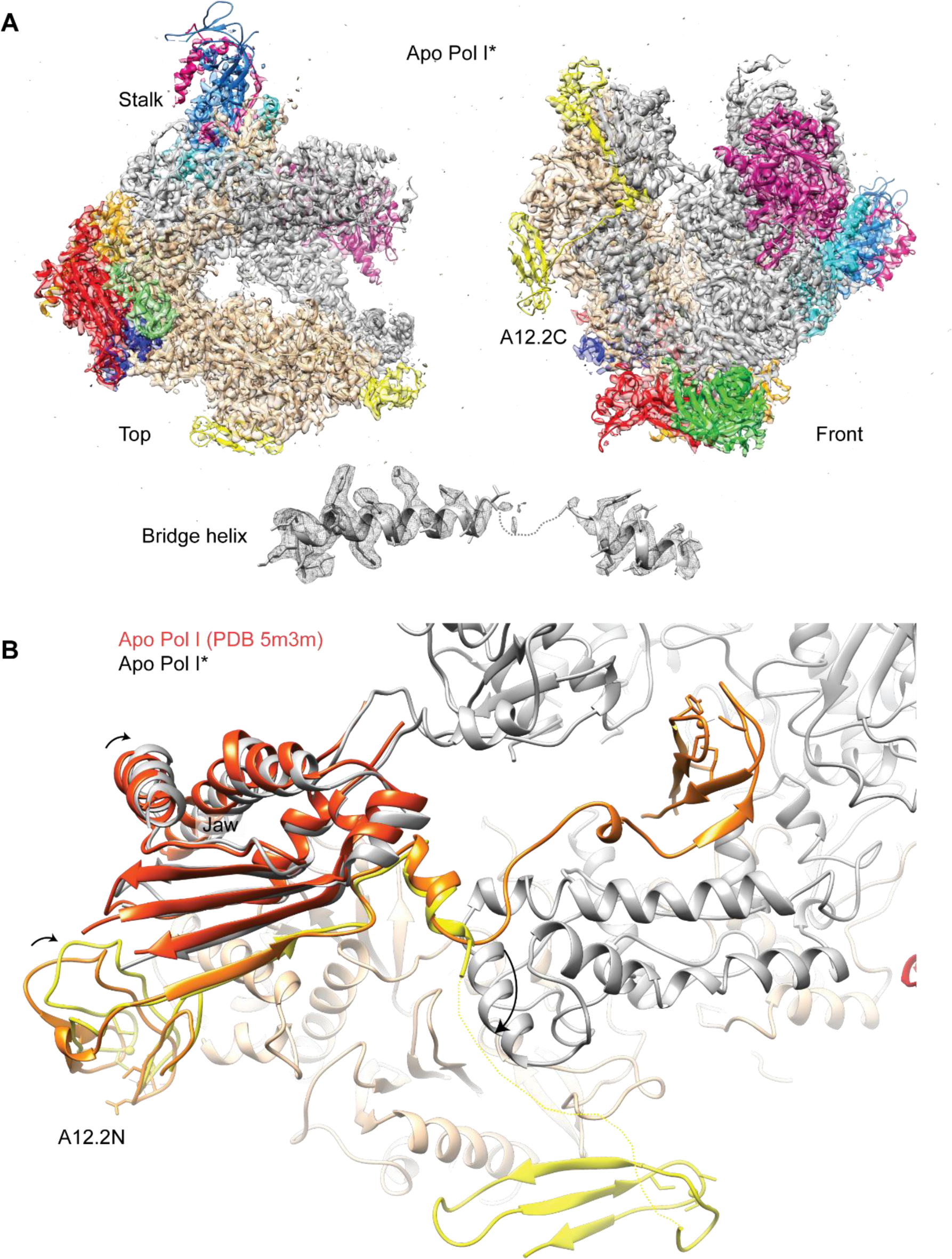
Structure of the apo Pol I*. **A.** Top and front views for the apo Pol I* model fitted into the sharpened density. The bridge helix is unfolded and shown below the models. **B.** Comparison between the A12.2 in the apo Pol I (PDB: 5m3m) and apo Pol I*. In the apo Pol I, the A12.2C is in the TFIIS-like position (orange), while in the apo Pol I*, the A12.2C is in the Rpb9-like position (yellow). Movement of the A12.2C induces a shift in the N-terminal A12.2 Zn ribbon domain A12.2N, which moves towards the lobe. The flexible jaw helices also move slightly compared to the apo Pol I (orange red).

**Figure 4-Figure Supplement 1.**
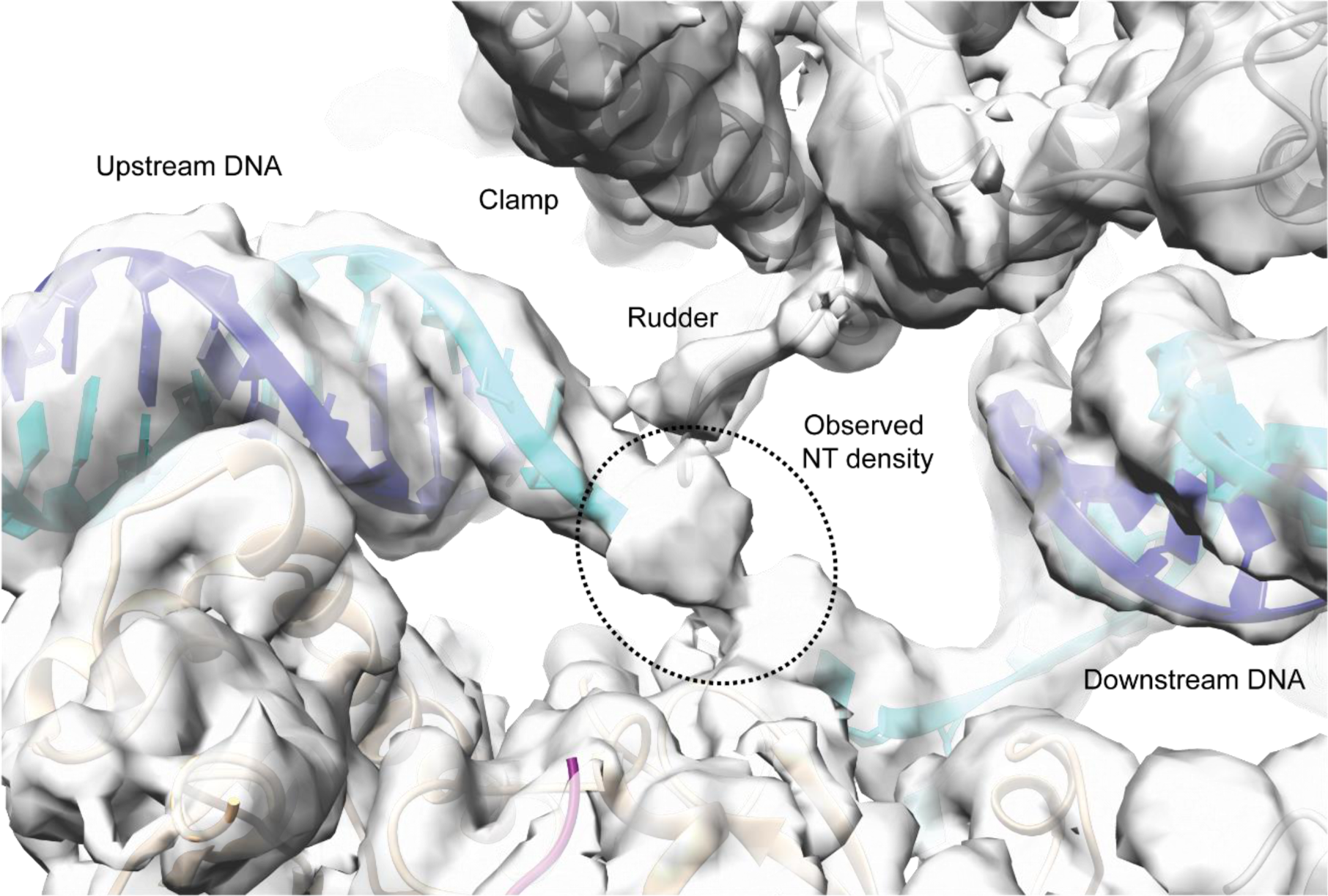
Density for the single-stranded non-template strand (ssNT) near the rudder. Density for the unsharpened and unmasked map for the Pol I EC focused on the upstream DNA with the fitted model shows that the density is bulky near the rudder.

**Figure 4-Figure Supplement 2.**
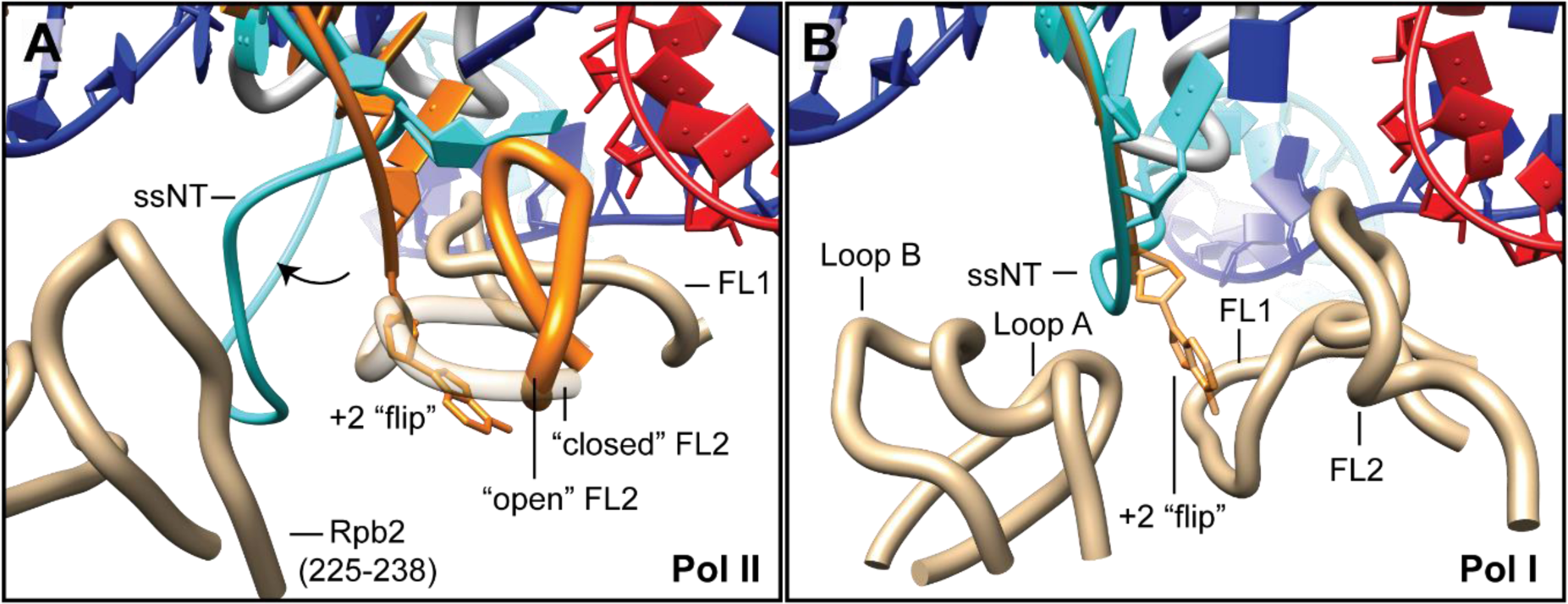
Conformations of the fork loop 2 in Pol II and Pol I. **A.** Two conformations of the Fork Loop 2 (FL2) are shown for Pol II, which affect the conformation of the +2 base. The FL2 open conformation which allows +2 base flipping(Cheung & Cramer, 2011) is shown in orange (PDB: 3po2), compared to the structure from the yeast Pol II EC with a complete transcription bubble (PDB: 5c4j). **B.** For comparison, the conformation of the downstream edge of the transcription bubble in Pol I is shown, in which the +2 base is flipped into the “A135 pocket”.

**Figure 4-Figure Supplement 3.**
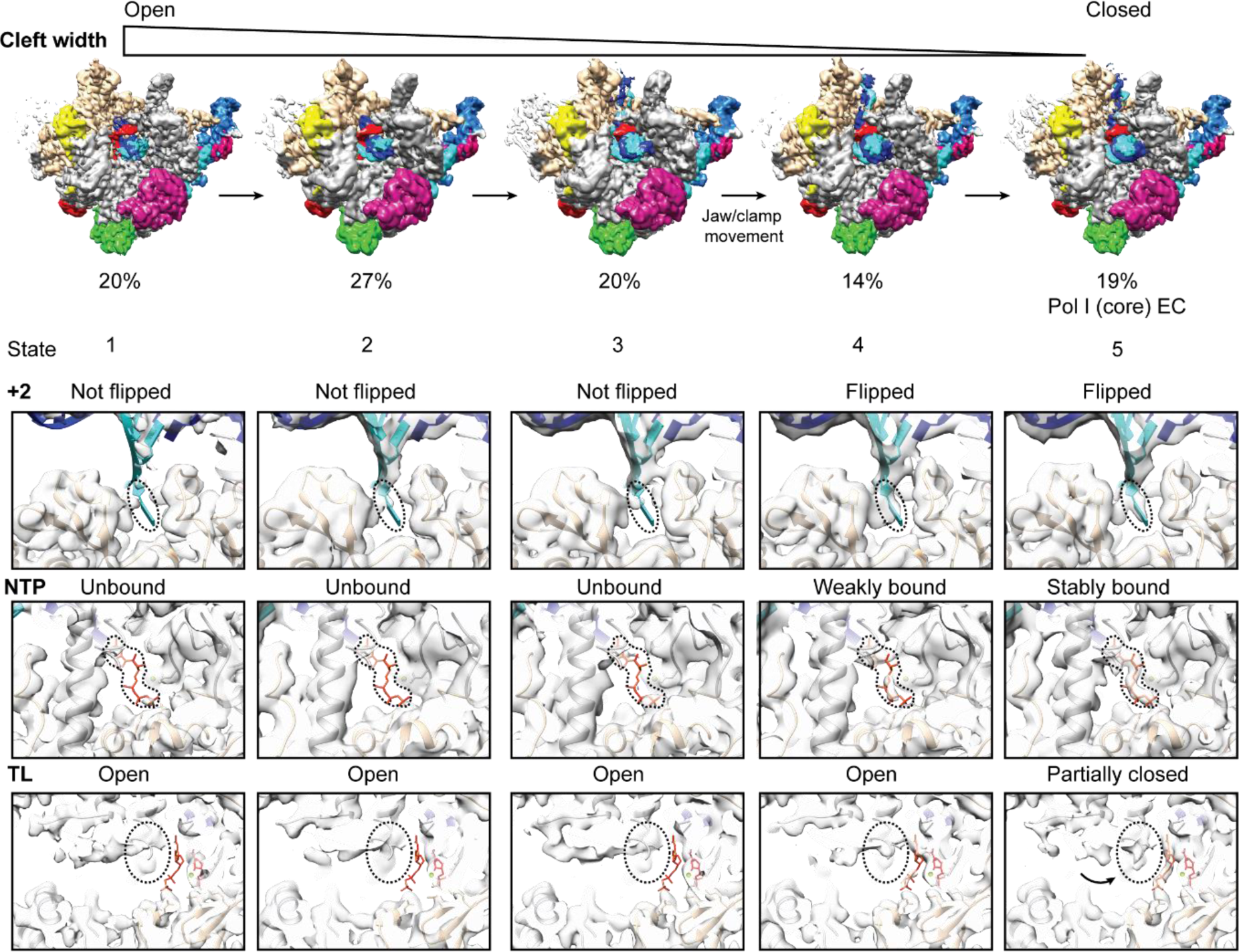
Conformational heterogeneity in the Pol I EC. Classification of the pooled Pol I EC particles reveals intermediates with slight differences in the width of the cleft, flipping of the +2 base, density for GMPCPP and trigger loop (TL). Closing of the cleft is accompanied by +2 base flipping, stronger density for GMPCPP and appearance of density for A190 L1202. Flipping of DNA base +2 occurs before complete closing of the cleft by the movement of the Jaw/clamp domains towards each other (between state 3 and 4). The transition from state 1 to 5 includes the sequential movement of the protrusion and wall domains towards the clamp, the jaw/clamp movement that induces +2 base flipping and closing of the cleft by movement of modules 1 and 2.

**Figure 5-Figure Supplement 1.**
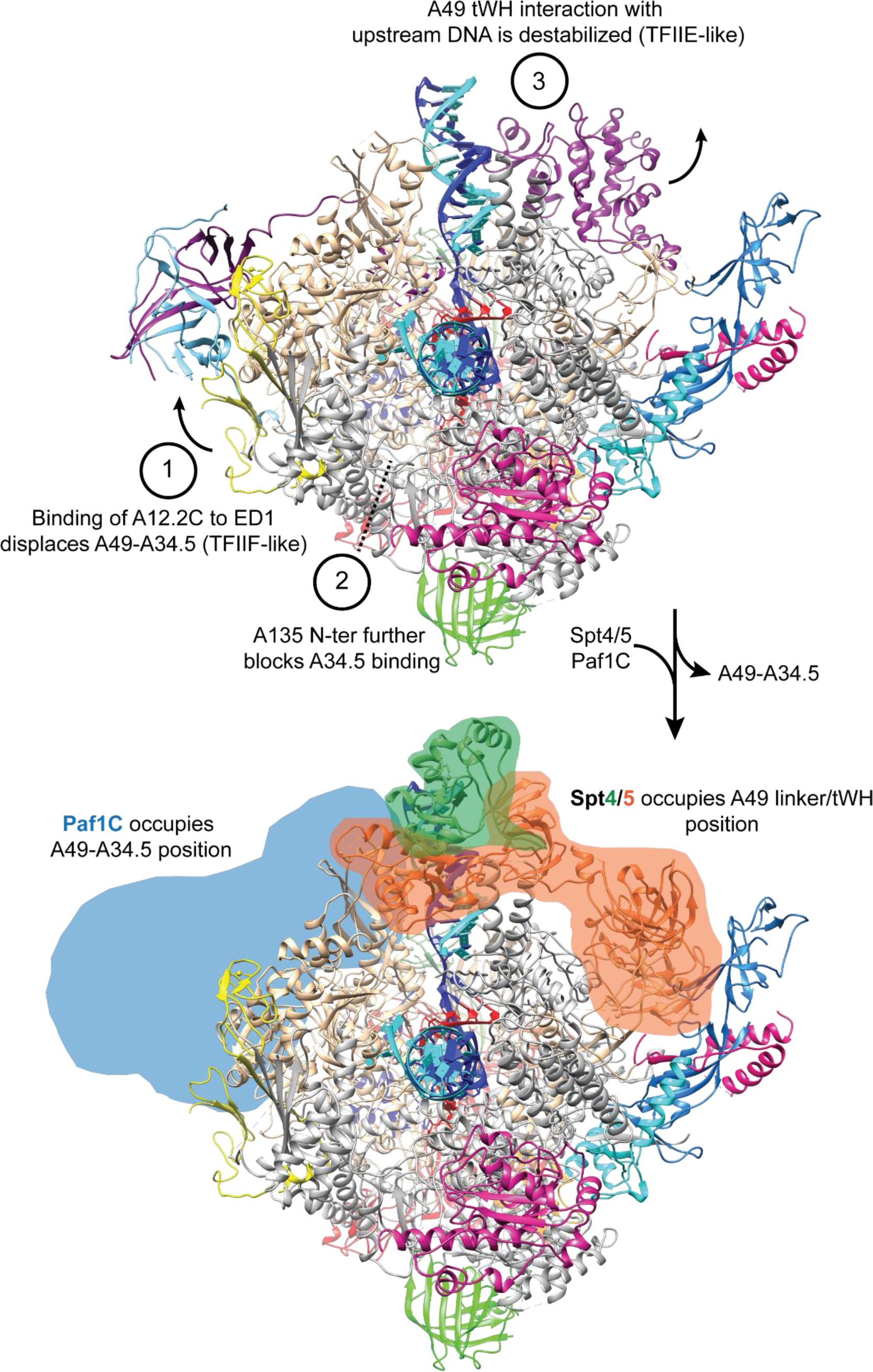
A49-A34.5 heterodimer release frees the binding site for Spt4/5 and Paf1C. Heterodimer binding to Pol I is destabilized by the binding of A12.2C to the ED1 (1) and binding of the A135-Nt to the HB further prevents interactions of Pol I with the A34.5-Ct (2). Destabilization of the interaction between the A49-A34.5 dimerization domain and Pol I removes the A49 tWH from the upstream DNA (3). Both A49-A34.5 dimerization module and A49 tWH sites are replaced by Paf1C and Spt4/5, respectively, in a manner similar to the transition from initiation to elongation in Pol II in which TFIIF and TFIIE are replaced by Paf1C and Spt4/5.

**Video 1. Jaw/clamp movement promote +2 base flipping.** +2 base flipping into the A135 pocket is induced by the relative movement of the jaw and clamp domains (transition from state 3 to 4 in Figure 4-Figure Supplement 3). A morph between maps corresponding to these states is shown.

**Video 2. Structural rearrangements upon Pol I activation.** The transition from state 1 to 5 in Figure 4-Figure Supplement 3 is shown by a morph of the maps corresponding to these states. This transition includes movement of the protrusion and wall, jaw and clamp, and complete closing of the cleft.

## References

Abascal-Palacios, G., Ramsay, E. P., Beuron, F., Morris, E., & Vannini, A. (2018). Structural basis of RNA polymerase III transcription initiation. Nature, 553(7688), 301–306. doi: 10.1038/nature25441

Adams, P. D., Afonine, P. V., Bunkoczi, G., Chen, V. B., Davis, I. W., Echols, N., … Zwart, P. H. (2010). PHENIX: a comprehensive Python-based system for macromolecular structure solution. Acta Crystallogr D Biol Crystallogr, 66(Pt 2), 213–221. doi: 10.1107/S0907444909052925

Albert, B., Leger-Silvestre, I., Normand, C., Ostermaier, M. K., Perez-Fernandez, J., Panov, K. I., … Gadal, O. (2011). RNA polymerase I-specific subunits promote polymerase clustering to enhance the rRNA gene transcription cycle. J Cell Biol, 192(2), 277–293. doi: 10.1083/jcb.201006040

Anderson, S. J., Sikes, M. L., Zhang, Y., French, S. L., Salgia, S., Beyer, A. L., … Schneider, D. A. (2011). The transcription elongation factor Spt5 influences transcription by RNA polymerase I positively and negatively. J Biol Chem, 286(21), 18816–18824. doi: 10.1074/jbc.M110.202101

Appling, F. D., Lucius, A. L., & Schneider, D. A. (2015). Transient-State Kinetic Analysis of the RNA Polymerase I Nucleotide Incorporation Mechanism. Biophys J, 109(11), 2382–2393. doi: 10.1016/j.bpj.2015.10.037

Appling, F. D., Schneider, D. A., & Lucius, A. L. (2017). Multisubunit RNA Polymerase Cleavage Factors Modulate the Kinetics and Energetics of Nucleotide Incorporation: An RNA Polymerase I Case Study. Biochemistry, 56(42), 5654–5662. doi: 10.1021/acs.biochem. 7b00370

Barnes, C. O., Calero, M., Malik, I., Graham, B. W., Spahr, H., Lin, G., … Calero, G. (2015). Crystal Structure of a Transcribing RNA Polymerase II Complex Reveals a Complete Transcription Bubble. Mol Cell, 59(2), 258–269. doi: 10.1016/j.molcel.2015.06.034

Beckouet, F., Mariotte-Labarre, S., Peyroche, G., Nogi, Y., & Thuriaux, P. (2011). Rpa43 and its partners in the yeast RNA polymerase I transcription complex. FEBS Lett, 585(21), 3355–3359. doi: 10.1016/j.febslet.2011.09.011

Bernecky, C., Herzog, F., Baumeister, W., Plitzko, J. M., & Cramer, P. (2016). Structure of transcribing mammalian RNA polymerase II. Nature, 529(7587), 551–554. doi: 10.1038/nature16482

Bernecky, C., Plitzko, J. M., & Cramer, P. (2017). Structure of a transcribing RNA polymerase II-DSIF complex reveals a multidentate DNA-RNA clamp. Nat Struct Mol Biol, 24(10), 809–815. doi: 10.1038/nsmb.3465

Cardone, G., Heymann, J. B., & Steven, A. C. (2013). One number does not fit all: mapping local variations in resolution in cryo-EM reconstructions. J Struct Biol, 184(2), 226–236. doi: 10.1016/j.jsb.2013.08.002

Chen, V. B., Arendall, W. B., 3rd, Headd, J. J., Keedy, D. A., Immormino, R. M., Kapral, G. J., … Richardson, D. C. (2010). MolProbity: all-atom structure validation for macromolecular crystallography. Acta Crystallogr D Biol Crystallogr, 66(Pt 1), 12–21. doi: 10.1107/S0907444909042073

Cheung, A. C., & Cramer, P. (2011). Structural basis of RNA polymerase II backtracking, arrest and reactivation. Nature, 471(7337), 249–253. doi: 10.1038/nature09785

Cheung, A. C., Sainsbury, S., & Cramer, P. (2011). Structural basis of initial RNA polymerase II transcription. EMBO J, 30(23), 4755–4763. doi: 10.1038/emboj.2011.396

Darrière T. P. M., Chauvier A, Genty T, Audibert S, Dez C, Leger-Silvestre I, Normand C, Calvo O, Fernández-Tornero C, Tschochner H, Gadal O. (2018). Genetic analysis of RNA polymerase I unveils new role of the Rpa12 subunit during transcription. bioRxiv. doi: https://doi.org/10.1101/307199

Ehara, H., Yokoyama, T., Shigematsu, H., Yokoyama, S., Shirouzu, M., & Sekine, S. I. (2017). Structure of the complete elongation complex of RNA polymerase II with basal factors. Science, 357(6354), 921–924. doi: 10.1126/science.aan8552

Emsley, P., & Cowtan, K. (2004). Coot: model-building tools for molecular graphics. Acta Crystallogr D Biol Crystallogr, 60(Pt 12 Pt 1), 2126–2132. doi: 10.1107/S0907444904019158

Engel, C., Gubbey, T., Neyer, S., Sainsbury, S., Oberthuer, C., Baejen, C., … Cramer, P. (2017). Structural Basis of RNA Polymerase I Transcription Initiation. Cell, 169(1), 120–131 e122. doi: 10.1016/j.cell.2017.03.003

Engel, C., Plitzko, J., & Cramer, P. (2016). RNA polymerase I-Rrn3 complex at 4.8 A resolution. Nat Commun, 7, 12129. doi: 10.1038/ncomms12129

Engel, C., Sainsbury, S., Cheung, A. C., Kostrewa, D., & Cramer, P. (2013). RNA polymerase I structure and transcription regulation. Nature, 502(7473), 650–655. doi: 10.1038/nature12712

Fernandez-Tornero, C., Moreno-Morcillo, M., Rashid, U. J., Taylor, N. M., Ruiz, F. M., Gruene, T., … Muller, C. W. (2013). Crystal structure of the 14-subunit RNA polymerase I. Nature, 502(7473), 644–649. doi: 10.1038/nature12636

Gadal, O., Mariotte-Labarre, S., Chedin, S., Quemeneur, E., Carles, C., Sentenac, A., & Thuriaux, P. (1997). A34.5, a nonessential component of yeast RNA polymerase I, cooperates with subunit A14 and DNA topoisomerase I to produce a functional rRNA synthesis machine. Mol Cell Biol, 17(4), 1787–1795.

Geiger, S. R., Lorenzen, K., Schreieck, A., Hanecker, P., Kostrewa, D., Heck, A. J., & Cramer, P. (2010). RNA polymerase I contains a TFIIF-related DNA-binding subcomplex. Mol Cell, 39(4), 583–594. doi: 10.1016/j.molcel.2010.07.028

Han, Y., Yan, C., Nguyen, T. H. D., Jackobel, A. J., Ivanov, I., Knutson, B. A., & He, Y. (2017). Structural mechanism of ATP-independent transcription initiation by RNA polymerase I. Elife, 6. doi: 10.7554/eLife.27414

He, Y., Yan, C., Fang, J., Inouye, C., Tjian, R., Ivanov, I., & Nogales, E. (2016). Near-atomic resolution visualization of human transcription promoter opening. Nature, 533(7603), 359–365. doi: 10.1038/nature17970

Hemming, S. A., Jansma, D. B., Macgregor, P. F., Goryachev, A., Friesen, J. D., & Edwards, A. M. (2000). RNA polymerase II subunit Rpb9 regulates transcription elongation in vivo. J Biol Chem, 275(45), 35506–35511. doi: 10.1074/jbc.M004721200

Hoffmann, N. A., Jakobi, A. J., Moreno-Morcillo, M., Glatt, S., Kosinski, J., Hagen, W. J., … Muller, C. W. (2015). Molecular structures of unbound and transcribing RNA polymerase III. Nature, 528(7581), 231–236. doi: 10.1038/nature16143

Huang, X., Wang, D., Weiss, D. R., Bushnell, D. A., Kornberg, R. D., & Levitt, M. (2010). RNA polymerase II trigger loop residues stabilize and position the incoming nucleotide triphosphate in transcription. Proc Natl Acad Sci U S A, 107(36), 15750–15745. doi: 10.1073/pnas.1009898107

Huet, J., Buhler, J. M., Sentenac, A., & Fromageot, P. (1975). Dissociation of two polypeptide chains from yeast RNA polymerase A. Proc Natl Acad Sci U S A, 72(8), 3034–3038.

Huet, J., Dezelee, S., Iborra, F., Buhler, J. M., Sentenac, A., & Fromageot, P. (1976). Further characterization of yeast RNA polymerases. Effect of subunits removal. Biochimie, 58(1-2), 71–80.

Kettenberger, H., Armache, K. J., & Cramer, P. (2004). Complete RNA polymerase II elongation complex structure and its interactions with NTP and TFIIS. Mol Cell, 16(6), 955–965. doi: 10.1016/j.molcel.2004.11.040

Khatter, H., Vorlander, M. K., & Muller, C. W. (2017). RNA polymerase I and III: similar yet unique. Curr Opin Struct Biol, 47, 88–94. doi: 10.1016/j.sbi.2017.05.008

Kimanius, D., Forsberg, B. O., Scheres, S. H., & Lindahl, E. (2016). Accelerated cryo-EM structure determination with parallelisation using GPUs in RELION-2. Elife, 5. doi: 10.7554/eLife.18722

Knippa, K., & Peterson, D. O. (2013). Fidelity of RNA polymerase II transcription: Role of Rpb9 [corrected] in error detection and proofreading. Biochemistry, 52(44), 7807–7817. doi: 10.1021/bi4009566

Kuhn, C. D., Geiger, S. R., Baumli, S., Gartmann, M., Gerber, J., Jennebach, S., … Cramer, P. (2007). Functional architecture of RNA polymerase I. Cell, 131(7), 1260–1272. doi: 10.1016/j.cell.2007.10.051

Li, S., Ding, B., Chen, R., Ruggiero, C., & Chen, X. (2006). Evidence that the transcription elongation function of Rpb9 is involved in transcription-coupled DNA repair in Saccharomyces cerevisiae. Mol Cell Biol, 26(24), 9430–9441. doi: 10.1128/MCB.01656-06

Liljelund, P., Mariotte, S., Buhler, J. M., & Sentenac, A. (1992). Characterization and mutagenesis of the gene encoding the A49 subunit of RNA polymerase A in Saccharomyces cerevisiae. Proc Natl Acad Sci U S A, 89(19), 9302–9305.

Lisica, A., Engel, C., Jahnel, M., Roldan, E., Galburt, E. A., Cramer, P., & Grill, S. W. (2016). Mechanisms of backtrack recovery by RNA polymerases I and II. Proc Natl Acad Sci U S A, 113(11), 2946–2951. doi: 10.1073/pnas.1517011113

Milkereit, P., & Tschochner, H. (1998). A specialized form of RNA polymerase I, essential for initiation and growth-dependent regulation of rRNA synthesis, is disrupted during transcription. EMBO J, 17(13), 3692–3703. doi: 10.1093/emboj/17.13.3692

Mishanina, T. V., Palo, M. Z., Nayak, D., Mooney, R. A., & Landick, R. (2017). Trigger loop of RNA polymerase is a positional, not acid-base, catalyst for both transcription and proofreading. Proc Natl Acad Sci U S A, 114(26), E5103–E5112. doi: 10.1073/pnas.1702383114

Moreno-Morcillo, M., Taylor, N. M., Gruene, T., Legrand, P., Rashid, U. J., Ruiz, F. M., … Fernandez-Tornero, C. (2014). Solving the RNA polymerase I structural puzzle. Acta Crystallogr D Biol Crystallogr, 70(Pt 10), 2570–2582. doi: 10.1107/S1399004714015788

Neyer, S., Kunz, M., Geiss, C., Hantsche, M., Hodirnau, V. V., Seybert, A., … Frangakis, A. S. (2016). Structure of RNA polymerase I transcribing ribosomal DNA genes. Nature. doi: 10.1038/nature20561

Pal, M., Ponticelli, A. S., & Luse, D. S. (2005). The role of the transcription bubble and TFIIB in promoter clearance by RNA polymerase II. Mol Cell, 19(1), 101–110. doi: 10.1016/j.molcel.2005.05.024

Penrod, Y., Rothblum, K., Cavanaugh, A., & Rothblum, L. I. (2015). Regulation of the association of the PAF53/PAF49 heterodimer with RNA polymerase I. Gene, 556(1), 61–67. doi: 10.1016/j.gene.2014.09.022

Petushkov, I., Pupov, D., Bass, I., & Kulbachinskiy, A. (2015). Mutations in the CRE pocket of bacterial RNA polymerase affect multiple steps of transcription. Nucleic Acids Res, 43(12), 5798–5809. doi: 10.1093/nar/gkv504

Pilsl, M., Crucifix, C., Papai, G., Krupp, F., Steinbauer, R., Griesenbeck, J., … Schultz, P.(2016). Structure of the initiation-competent RNA polymerase I and its implication for transcription. Nat Commun, 7, 12126. doi: 10.1038/ncomms12126

Plaschka, C., Hantsche, M., Dienemann, C., Burzinski, C., Plitzko, J., & Cramer, P. (2016). Transcription initiation complex structures elucidate DNA opening. Nature, 533(7603), 353–358. doi: 10.1038/nature17990

Rohou, A., & Grigorieff, N. (2015). CTFFIND4: Fast and accurate defocus estimation from electron micrographs. J Struct Biol, 192(2), 216–221. doi: 10.1016/j.jsb.2015.08.008

Rossbach, S., & Ochsenfeld, C. (2017). Quantum-Chemical Study of the Discrimination against dNTP in the Nucleotide Addition Reaction in the Active Site of RNA Polymerase II. J Chem Theory Comput, 13(4), 1699–1705. doi: 10.1021/acs.jctc.7b00157

Sadian, Y., Tafur, L., Kosinski, J., Jakobi, A. J., Wetzel, R., Buczak, K., … Muller, C. W. (2017) Structural insights into transcription initiation by yeast RNA polymerase I. EMBO J, 36(18), 2698–2709. doi: 10.15252/embj.201796958

Scheres, S. H. (2016). Processing of Structurally Heterogeneous Cryo-EM Data in RELION. Methods Enzymol, 579, 125–157. doi: 10.1016/bs.mie.2016.04.012

Schneider, D. A., Michel, A., Sikes, M. L., Vu, L., Dodd, J. A., Salgia, S., … Nomura, M. (2007). Transcription elongation by RNA polymerase I is linked to efficient rRNA processing and ribosome assembly. Mol Cell, 26(2), 217–229. doi: 10.1016/j.molcel.2007.04.007

Tafur, L., Sadian, Y., Hoffmann, N. A., Jakobi, A. J., Wetzel, R., Hagen, W. J. H., … Muller, C. W. (2016). Molecular Structures of Transcribing RNA Polymerase I. Mol Cell, 64(6), 1135–1143. doi: 10.1016/j.molcel.2016.11.013

Torreira, E., Louro, J. A., Pazos, I., Gonzalez-Polo, N., Gil-Carton, D., Duran, A. G., …Fernandez-Tornero, C. (2017). The dynamic assembly of distinct RNA polymerase I complexes modulates rDNA transcription. Elife, 6. doi: 10.7554/eLife.20832

Van Mullem, V., Landrieux, E., Vandenhaute, J., & Thuriaux, P. (2002). Rpa12p, a conserved RNA polymerase I subunit with two functional domains. Mol Microbiol, 43(5), 1105–1113.

Vannini, A., & Cramer, P. (2012). Conservation between the RNA polymerase I, II, and III transcription initiation machineries. Mol Cell, 45(4), 439–446. doi: 10.1016/j.molcel.2012.01.023

Vassylyev, D. G., Vassylyeva, M. N., Zhang, J., Palangat, M., Artsimovitch, I., & Landick, R. (2007). Structural basis for substrate loading in bacterial RNA polymerase. Nature, 448(7150), 163–168. doi: 10.1038/nature05931

Viktorovskaya, O. V., Appling, F. D., & Schneider, D. A. (2011). Yeast transcription elongation factor Spt5 associates with RNA polymerase I and RNA polymerase II directly. J Biol Chem, 286(21), 18825–18833. doi: 10.1074/jbc.M110.202119

Viktorovskaya, O. V., Engel, K. L., French, S. L., Cui, P., Vandeventer, P. J., Pavlovic, E. M., … Schneider, D. A. (2013). Divergent contributions of conserved active site residues to transcription by eukaryotic RNA polymerases I and II. Cell Rep, 4(5), 974–984. doi: 10.1016/j.celrep.2013.07.044

Vorlander, M. K., Khatter, H., Wetzel, R., Hagen, W. J. H., & Muller, C. W. (2018). Molecular mechanism of promoter opening by RNA polymerase III. Nature, 553(7688), 295–300. doi: 10.1038/nature25440

Vvedenskaya, I. O., Vahedian-Movahed, H., Bird, J. G., Knoblauch, J. G., Goldman, S. R., Zhang, Y., … Nickels, B. E. (2014). Interactions between RNA polymerase and the “core recognition element” counteract pausing. Science, 344(6189), 1285–1289. doi: 10.1126/science.1253458

Vvedenskaya, I. O., Vahedian-Movahed, H., Zhang, Y., Taylor, D. M., Ebright, R. H., & Nickels, B. E. (2016). Interactions between RNA polymerase and the core recognition element are a determinant of transcription start site selection. Proc Natl Acad Sci U S A, 113(21), E2899–2905. doi: 10.1073/pnas.1603271113

Wang, D., Bushnell, D. A., Westover, K. D., Kaplan, C. D., & Kornberg, R. D. (2006). Structural basis of transcription: role of the trigger loop in substrate specificity and catalysis. Cell, 127(5), 941–954. doi: 10.1016/j.cell.2006.11.023

Westover, K. D., Bushnell, D. A., & Kornberg, R. D. (2004). Structural basis of transcription: nucleotide selection by rotation in the RNA polymerase II active center. Cell, 119(4), 481–489. doi: 10.1016/j.cell.2004.10.016

Wiesler, S. C., Weinzierl, R. O., & Buck, M. (2013). An aromatic residue switch in enhancer-dependent bacterial RNA polymerase controls transcription intermediate complex activity. Nucleic Acids Res, 41(11), 5874–5886. doi: 10.1093/nar/gkt271

Xu, Y., Bernecky, C., Lee, C. T., Maier, K. C., Schwalb, B., Tegunov, D., … Cramer, P. (2017). Architecture of the RNA polymerase I I-Paf1 C-TFIIS transcription elongation complex. Nat Commun, 8, 15741. doi: 10.1038/ncomms15741

Yamamoto, K., Yamamoto, M., Hanada, K., Nogi, Y., Matsuyama, T., & Muramatsu, M. (2004). Multiple protein-protein interactions by RNA polymerase I-associated factor PAF49 and role of PAF49 in rRNA transcription. Mol Cell Biol, 24(14), 6338–6349. doi: 10.1128/MCB.24.14.6338-6349.2004

Zhang, Y., Feng, Y., Chatterjee, S., Tuske, S., Ho, M. X., Arnold, E., & Ebright, R. H. (2012). Structural basis of transcription initiation. Science, 338(6110), 1076–1080. doi: 10.1126/science.1227786

Zhang, Y., Sikes, M. L., Beyer, A. L., & Schneider, D. A. (2009). The Paf1 complex is required for efficient transcription elongation by RNA polymerase I. Proc Natl Acad Sci U S A, 106(7), 2153–2158. doi: 10.1073/pnas.0812939106

Zhang, Y., Smith, A. D. t., Renfrow, M. B., & Schneider, D. A. (2010). The RNA polymerase-associated factor 1 complex (Paf1C) directly increases the elongation rate of RNA polymerase I and is required for efficient regulation of rRNA synthesis. J Biol Chem, 285(19), 14152–14159. doi: 10.1074/jbc.M110.115220

Zheng, S. Q., Palovcak, E., Armache, J. P., Verba, K. A., Cheng, Y., & Agard, D. A. (2017). MotionCor2: anisotropic correction of beam-induced motion for improved cryo-electron microscopy. Nat Methods, 14(4), 331–332. doi: 10.1038/nmeth.4193

